# Striatal Dopamine at Learned Sequence Boundaries Sustains Birdsong

**DOI:** 10.64898/2026.07.05.736606

**Authors:** Lei Xiao, Todd F. Roberts

**Affiliations:** Department of Neuroscience, UT Southwestern Medical Center, Dallas TX, USA

## Abstract

Phasic striatal dopamine has been implicated in the initiation of well-trained action sequences in reward-guided tasks. Whether such signals also support natural skills learned without explicit cues or immediate rewards remains unknown. Using birdsong as a model of a naturally learned, skilled vocal behavior, we found that dopamine transients accompany the initiation of song sequences. These transients emerged during learning, shifting from later phases of the sequence toward sequence onset as song matured. Temporally targeted optogenetic inhibition of dopamine signaling at sequence onsets disrupted the maintenance of learned song, resulting in the gradual and severe deterioration of adult song. Thus, phasic dopamine signaling at sequence initiation develops during learning of a natural skilled behavior and is required for its long-term maintenance.

**One Sentence Summary:** Dopamine transients at song motif onset are required to maintain accurate adult zebra finch song, a naturally learned skilled behavior.

## Main

Skilled behaviors require the brain to initiate organized action sequences rather than isolated movements. Dorsal striatal circuits have been implicated in this process(*1–5*). Neural activity in the striatum marks the initiation and termination of learned sequences, suggesting a role in defining the boundaries of higher-order behavioral units(*5–7*).

Phasic dopamine release is a candidate signal for gating these sequence boundaries. Although striatal dopamine transients are classically associated with reward prediction errors(*8–10*), they also accompany self-initiated actions(*11–14*) and can precede the onset of well-trained operant sequences(*6, 15, 16*). These findings raise the possibility that dopamine helps launch learned action programs. Yet, most evidence linking dopamine to sequence initiation comes from tasks built around explicit rewards, discrete cues, and stereotyped responses. Natural skills differ from such operant tasks. They emerge through extended practice, rely on internal performance feedback, and persist long after learning. It therefore remains unclear whether dopamine transients organize naturally acquired skilled behaviors or instead reflect reward-trained task structure.

Birdsong provides a tractable model for addressing this question. Zebra finch song is learned during development and produced in adulthood without external instruction or immediate reward. Its discrete vocal gestures, or syllables, are organized into a stereotyped repeated unit, the song motif. In zebra finches, dopamine activity in the song basal ganglia(sBG) has been shown to track vocal performance feedback and the quality of individual syllables during learning(*17–23*), supporting a syllable-level performance-evaluation and error-correction model(*24–26*). Whether dopamine is also organized at a higher level, around motif boundaries, and whether such signals support song maintenance in adulthood, remains unknown.

### Striatal Dopamine Peaks at Motif Initiation in Adult Song

Prior electrophysiological recordings from the ventral tegmental area (VTA) in juvenile zebra finches have revealed increased activity near song-bout onset(*27*), suggesting that dopamine neurons may be recruited during vocal sequence initiation. This motivated us to ask whether dopamine release in the sBG is also temporally organized around individual song motifs. To address this question, we expressed the dopamine sensor dLight1.3b in Area X, a striatal nucleus of the sBG, and used fiber photometry to monitor dopamine-dependent fluorescence during adult singing (**Fig. 1, A and B**). This approach revealed rapid dopamine fluctuations during song motifs, which had an average duration of 1,015 ± 25 ms.

**Figure 1.**
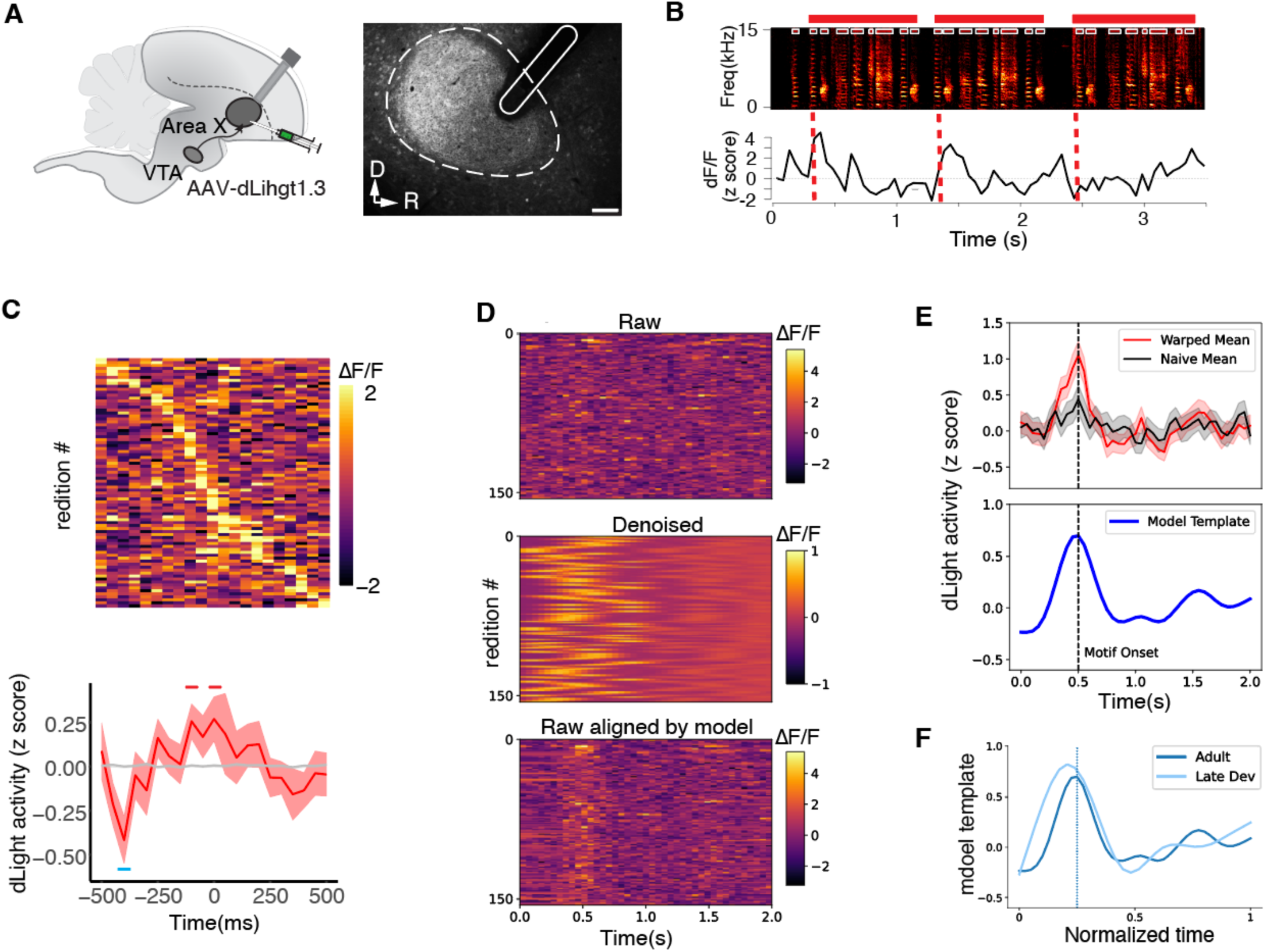
Striatal Dopamine Peaks at Motif Initiation in Adult Song. **A)** Left, schematic of AAV-dLight1.3 injection into Area X and fiber placement over Area X. Right, representative dLight1.3 expression in Area X. Dashed outline marks Area X; white outline indicates fiber track. Scale bar, 250 μm. **B)** Example spectrogram and simultaneously recorded standardized dLight fluorescence during singing. Red dashed lines mark motif onsets; red bars indicate motif renditions. **C)** Trial-by-trial dLight fluorescence during directed singing, aligned to motif onset and sorted by peak timing. Bottom, median motif-onset-aligned dLight activity. Red bars indicate significant positive deviations from shuffle controls near motif onset (−100 ms, p = 0.011; 0 ms, p = 0.011; other time points, p > 0.05, shuffle test), and the blue bar indicates a significant dip within the motif (−400 ms, p = 0.019). Grey shading indicates the 95% shuffle confidence interval (CI); shaded error denotes ±s.e.m. **D)** Trial-by-trial dLight fluorescence during singing shown as raw signals, denoised signals, and signals realigned by the shift-only model. **E)** Top, average dLight activity across motif renditions using naive alignment (black) or shift-only time-warping alignment (red). Shading indicates 95% CI. Bottom, model-derived template of dLight activity showing a consolidated peak at motif onset and a smaller rebound near motif offset. Dashed line marks motif onset. **F)** Model-derived dLight templates from adult and late-development song, plotted in normalized motif time. Both show dopamine activity concentrated near motif onset.

We first manually segmented motifs from female-directed song and aligned dopamine signals to the onset or offset of motifs (**fig. S1A**). Motif-onset alignment revealed a stereotyped dip-then-peak pattern: dopamine fluorescence decreased several hundred milliseconds before motif onset, peaked near motif initiation, and declined as the motif unfolded (**Fig. 1C**). This temporal profile resembles reported burst-pause dynamics in dopamine neurons during movement initiation(*11, 13, 28*). Motif-offset alignment also revealed a preceding dip, although the post-offset peak was less distinct (**fig. S1B**). Around motif onset, dopamine signals were correlated across hemispheres, and detected transients became more frequent, indicating coordinated striatal dopamine activity at motif initiation (**fig. S1, C to F**).

To test whether this motif-onset peak depended on manual alignment, we applied an unsupervised shift-only time-warping model(*29*) to dopamine traces around each motif (**Fig. 1D**). This procedure realigned trials by temporal offset, without stretching or compressing the fluorescence traces. In pooled directed and undirected song renditions, this data-driven alignment revealed a consistent dopamine peak at motif onsets, followed by a smaller rebound near motif offset (**Fig. 1E**). The same temporal pattern was observed in both continuous fluorescence signals and detected transients (**fig. S2, A to C**), indicating that motif-locked dopamine dynamics are not an artifact of manual behavioral alignment.

### Dopamine Signals Shift from Motif Offset to Onset During Song Development

We next asked whether ‘motif-onset’ dopamine signals emerge during developmental learning. Recent longitudinal studies in zebra finches showed that dopamine activity in the sBG tracks the performance quality of individual syllables during song acquisition(21, 22). Reanalysis of publicly available data from Qi et al. revealed a similar but broader GRAB-DA signal at motif onset late in development (89 ± 1 dph, days post hatch), when song approaches crystallization (**Fig. 1F; fig. S3, A and B**). This signal was not evident earlier in development (70 ± 1 dph; **fig. S3C**), suggesting that motif-onset dopamine activity emerges as song is consolidated into a stable learned action sequence.

These observations raised the question of whether motif-onset dopamine in adult song is analogous to cue-locked dopamine in well-trained reward-guided behaviors. In laboratory training paradigms, dopamine activity often peaks early in learning at reward delivery or sequence completion, but shifts with learning toward earlier predictive events, including cues or action onsets(*8, 15, 16, 30–32*). This pattern is commonly interpreted as being consistent with temporal-difference (TD) learning(*33, 34*). Song, however, is acquired without discrete external cues or immediate rewards, making it unclear whether a similar predictive dopamine organization can emerge during natural skill learning.

The developmental data offered a clue. Although dopamine did not peak at motif onset early in development, time-warping analysis revealed a lower-amplitude peak near motif offset (**fig. S3, A to C**). If motif-onset dopamine in adult song arises through a TD-like developmental process, song learning should be associated with a backward shift in dopamine peak timing, redistribution of dopamine amplitude from later to earlier motif segments, and the emergence of anticipatory relationships between early and late motif segment activity (**Fig. 2A**).

**Figure 2.**
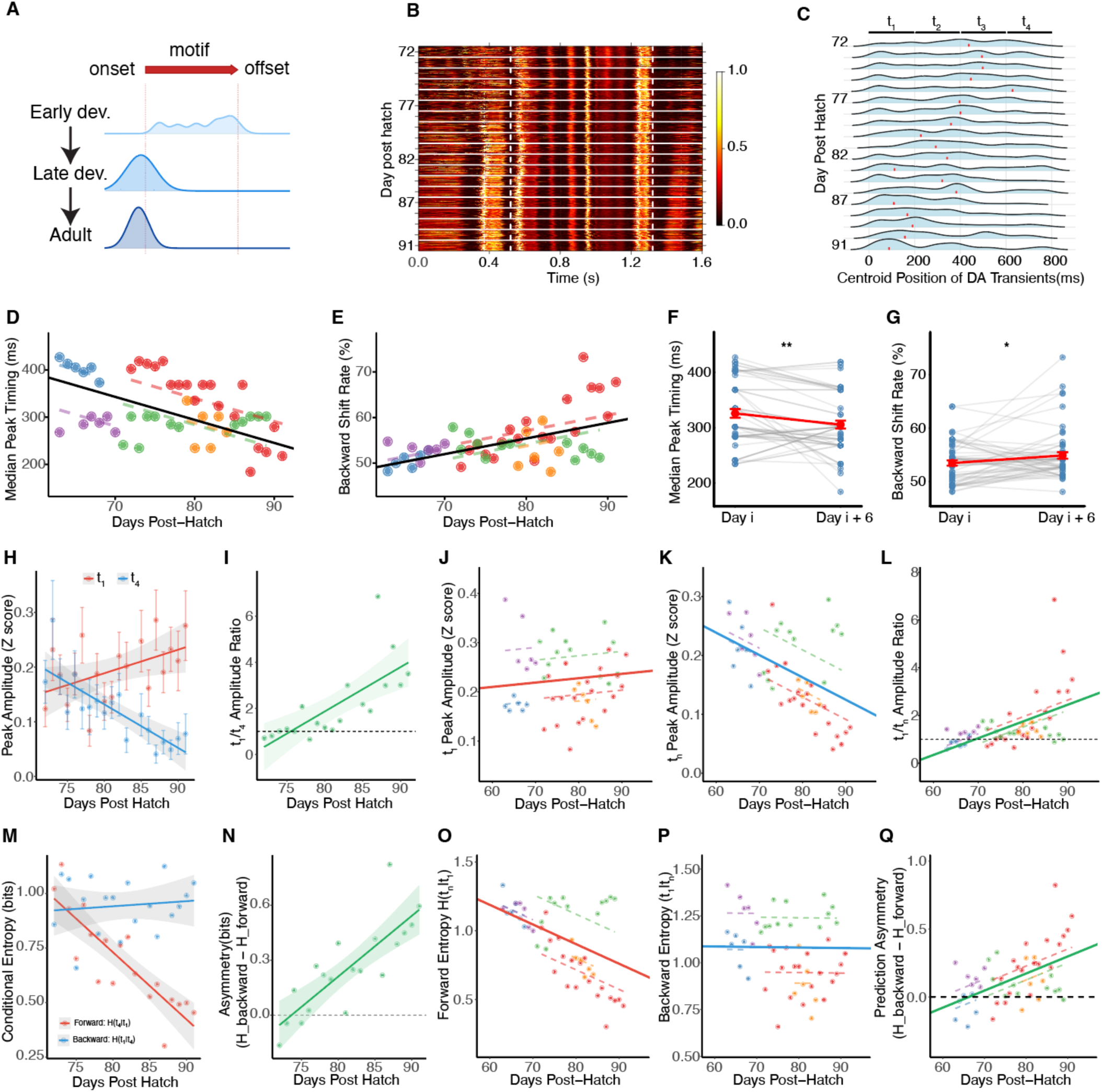
Developmental Reorganization of Dopamine Activity During Vocal Learning. **A)** Schematic illustrating the proposed developmental reorganization of dopamine activity across the song motif. **B)** Amplitude envelopes of aligned song motifs from an example bird across development. Each row represents one epoch; dashed lines mark motif onset and offset. **C)** Density ridge plots showing the developmental distribution of dopamine-transient centroid positions in the example bird. Red ticks mark distribution peaks; t_1_-t_4_ indicate 200-ms motif segments. **D)** – **G**) Population developmental trajectories of dopamine-transient timing metrics. Median peak timing decreased with age (**D**), whereas backward shift rate increased (**E**). Six-day sliding-window analyses showed a corresponding decrease in peak timing (**F**) and increase in backward shift rate (**G**). **H)** – **L**) Developmental redistribution of transient amplitude. In an example bird, early-segment (t_1_) amplitude increased and late-segment (t_4_) amplitude decreased (**H**), producing an increased t_1_/t_4_ amplitude ratio (**I**). Across birds, early amplitude changed modestly (**J**), late amplitude decreased (**K**), and the t_1_/t_4_ ratio increased with age (**L**). **M)** – **Q**) Development of directional predictability across the motif. In an example bird, forward conditional entropy H(t_4_|t_1_) decreased while backward conditional entropy H(t_1_|t_4_) remained stable (**M**), producing increased prediction asymmetry (**N**). Across birds, forward conditional entropy decreased (**O**), backward conditional entropy showed little developmental change (**P**), and prediction asymmetry increased with age (**Q**). Statistics are summarized in Table 1.

**Table 1.**
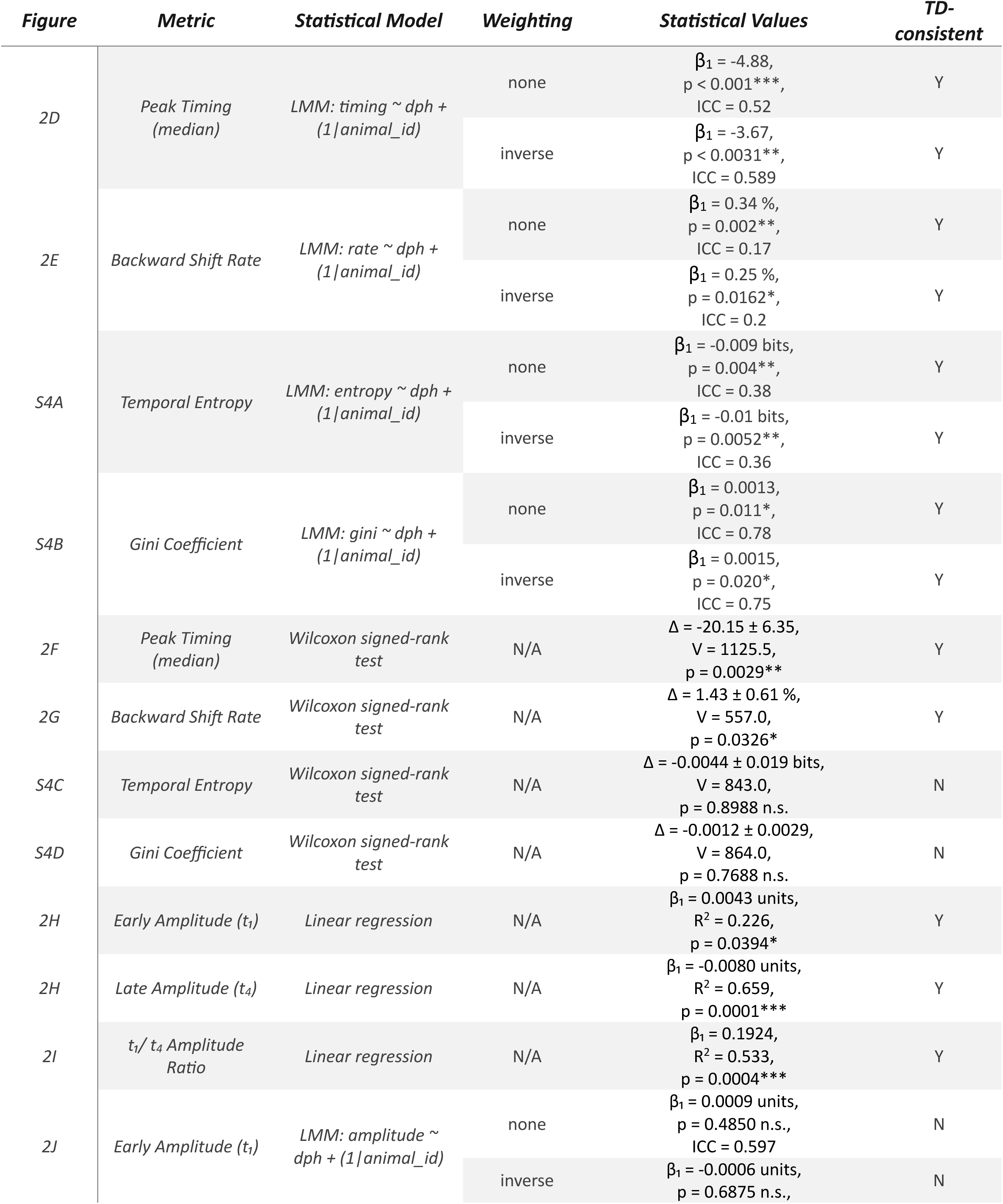
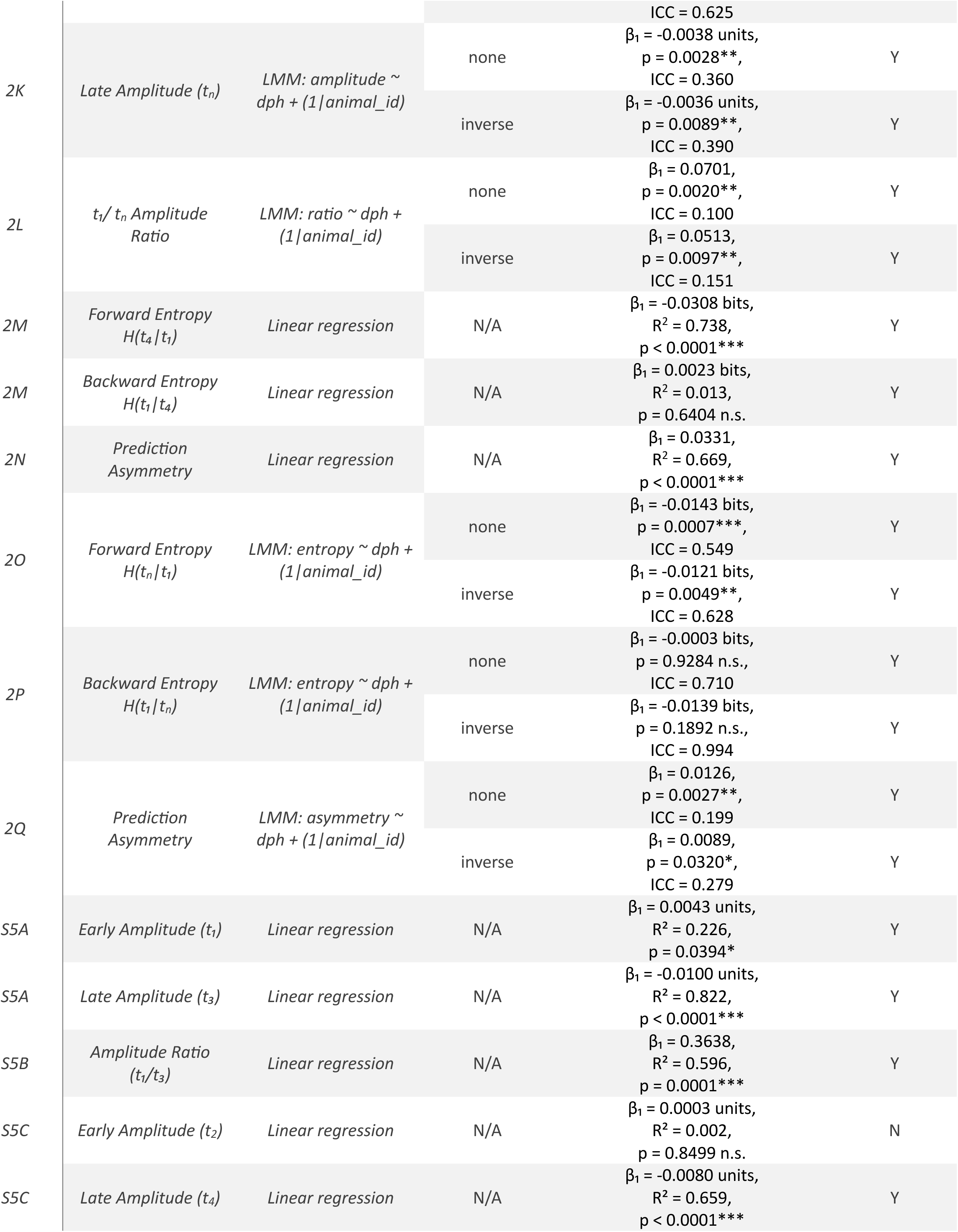
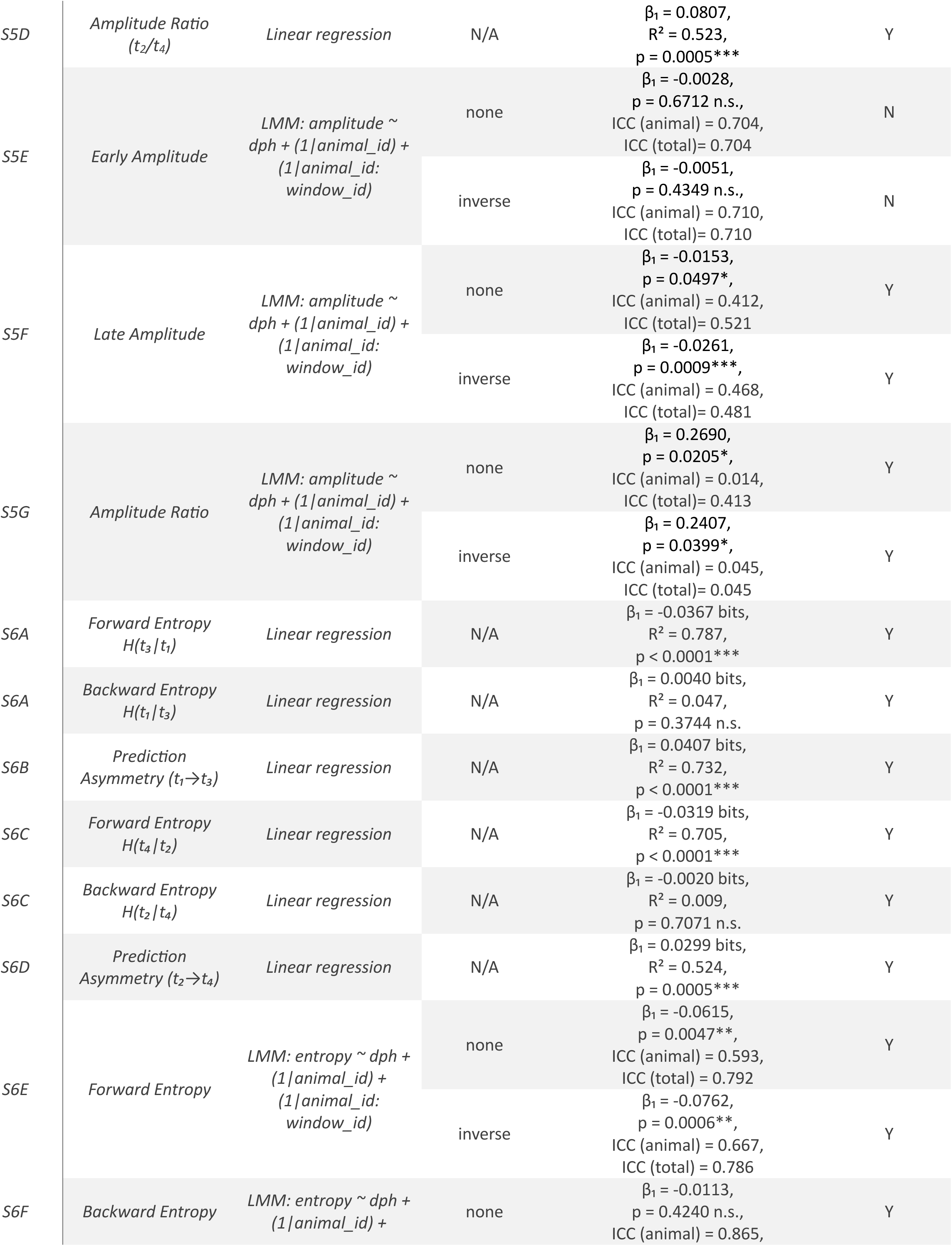
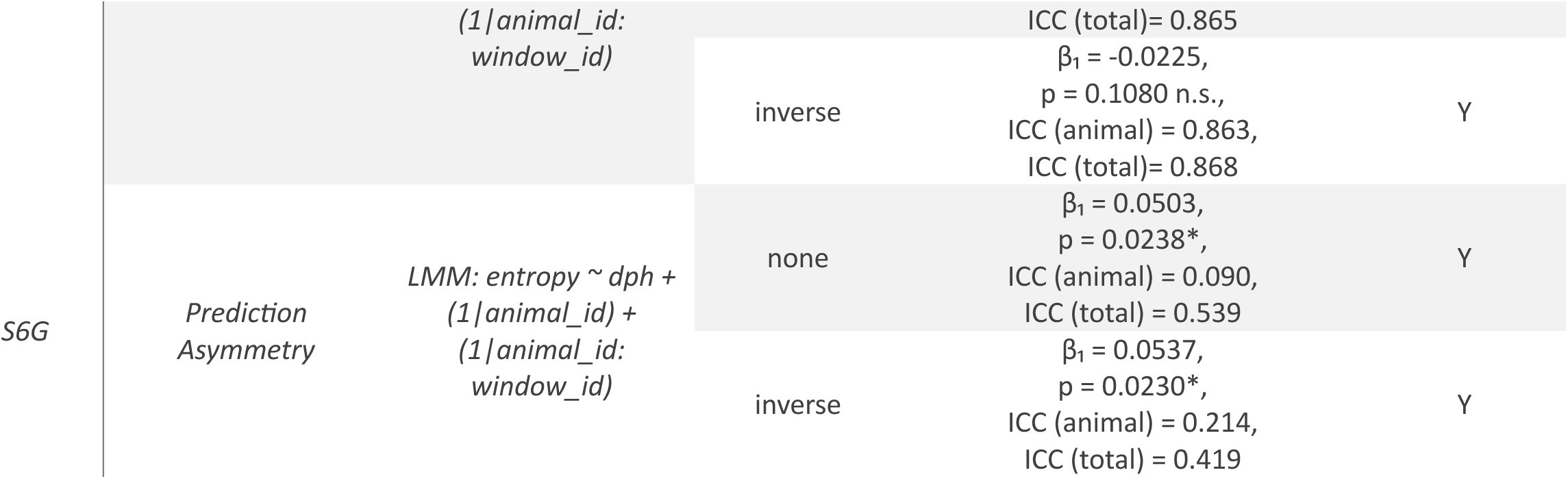
Statistical analyses of developmental fiber photometry metrics (Fig. 2; fig. S4-6)

To test these predictions, we analyzed motif-level dopamine activity from 7,029 motifs collected across 433 recording sessions in five juvenile birds between 63 and 91 dph. This longitudinal sampling allowed us to follow the reorganization of motif-related dopamine activity across vocal development. Using reliably anchored motif windows (**Fig. 2B**; see Methods), we quantified three features of this reorganization: peak timing, onset-to-offset amplitude ratios, and directional predictability between motif segments. Together, these analyses characterized how a broad, offset-biased signal in early development is reorganized toward the consolidated motif-onset signal observed in late development and adulthood.

Across development, peak dopamine activity shifted progressively from motif offset toward motif onset. In a representative bird, dopamine activity moved from later to earlier motif segments as song became more stereotyped (**Fig. 2C**). Across birds, linear mixed-effects models revealed a significant decrease in median peak time and a corresponding increase in a backward-shift index, which quantified redistribution of activity from motif offset toward onset (**Fig. 2, D and E**). These effects were robust to inverse sample-size weighting and balanced bootstrap resampling, indicating that they were not driven by unequal sampling across birds (**Table 1**; see Methods). Thus, dopamine activity became progressively aligned with motif initiation during vocal learning.

This developmental shift was accompanied by temporal sharpening. Entropy decreased with age, whereas the Gini coefficient increased, indicating that dopamine activity became concentrated within narrower motif windows (**Fig. S4, A and B**). Six-day developmental sliding-window analyses showed that peak timing shifted earlier even over short developmental intervals, whereas entropy and Gini changed more gradually (**Fig. 2, F and G; fig. S4, C and D**). Thus, the temporal position and temporal precision of dopamine transients matured on partly distinct timescales.

Dopamine transient amplitude was also redistributed across the motif. Within individual birds, early-segment amplitudes increased whereas late-segment amplitudes decreased, producing a marked rise in the early-to-late amplitude ratio (**Fig. 2, H and I; fig. S5, A to D**). Across birds, the age-dependent increase in this ratio was driven most consistently by declining late-segment responses (**Fig. 2, J to L**). Six-day developmental sliding-window analyses across additional segment pairs separated by 200 ms revealed that amplitude redistribution occurred broadly across the motif and was detectable over short developmental intervals. (**fig. S5, A to G**).

Finally, we asked whether dopamine activity developed anticipatory structure across motif segments. Using directional conditional entropy, we tested whether activity in early segments increasingly predicted activity in later segments over development. Forward conditional entropy, which measures uncertainty about later activity given earlier activity, declined significantly with age, whereas backward conditional entropy remained relatively stable, producing an increasing forward-predictive asymmetry that emerged around 66 dph (**Fig. 2, M to Q**). The same forward-predictive pattern was evident in six-day developmental sliding-window analyses across additional segment pairs, indicating that anticipatory relationships between motif segments strengthened locally during development (**fig. S6, A to G**).

Together, these analyses show that dopamine activity is progressively shifted, sharpened, and reweighted toward motif onset during vocal learning, while acquiring forward-predictive structure across the learned sequence. These dynamics are consistent with a TD-like process in which dopamine signaling is progressively reorganized from an internally defined evaluative boundary at motif completion toward motif onset, an earlier predictive feature of the learned sequence.

### Phasic Inhibition of Dopamine Signaling at Motif Transitions Induces Song Decrystallization

The developmental reorganization of dopamine signaling toward motif onset suggested that this signal is linked to the consolidation of song as a stable action sequence. This raised a functional question: does motif-onset dopamine simply represent a residual trace of learning, or is it important for preserving the learned sequence following developmental learning? This distinction is important because dopamine signaling in the sBG has largely been framed as providing syllable-level performance feedback that guides adaptive vocal plasticity(24–26). If the motif-onset transient is functionally required in adults, it would indicate that dopamine also operates at a higher hierarchical level in the control of motor sequences, a previously unrecognized role for dopamine in natural behavior. We therefore asked whether temporally targeted perturbation of dopamine signaling near motif transitions would alter the structure or stability of adult song.

To suppress dopamine release with temporal precision, we bilaterally expressed the axon-targeted proton pump nxArchT in VTA dopamine neurons and implanted optical fibers over their terminals in Area X(*19*). Using closed-loop song detection, we delivered 100-ms optical inhibition during singing (**Fig. 3A**). The inhibition window was fixed within each bird but varied across birds, allowing us to sample perturbations delivered at different times relative to the song motif.

**Figure 3.**
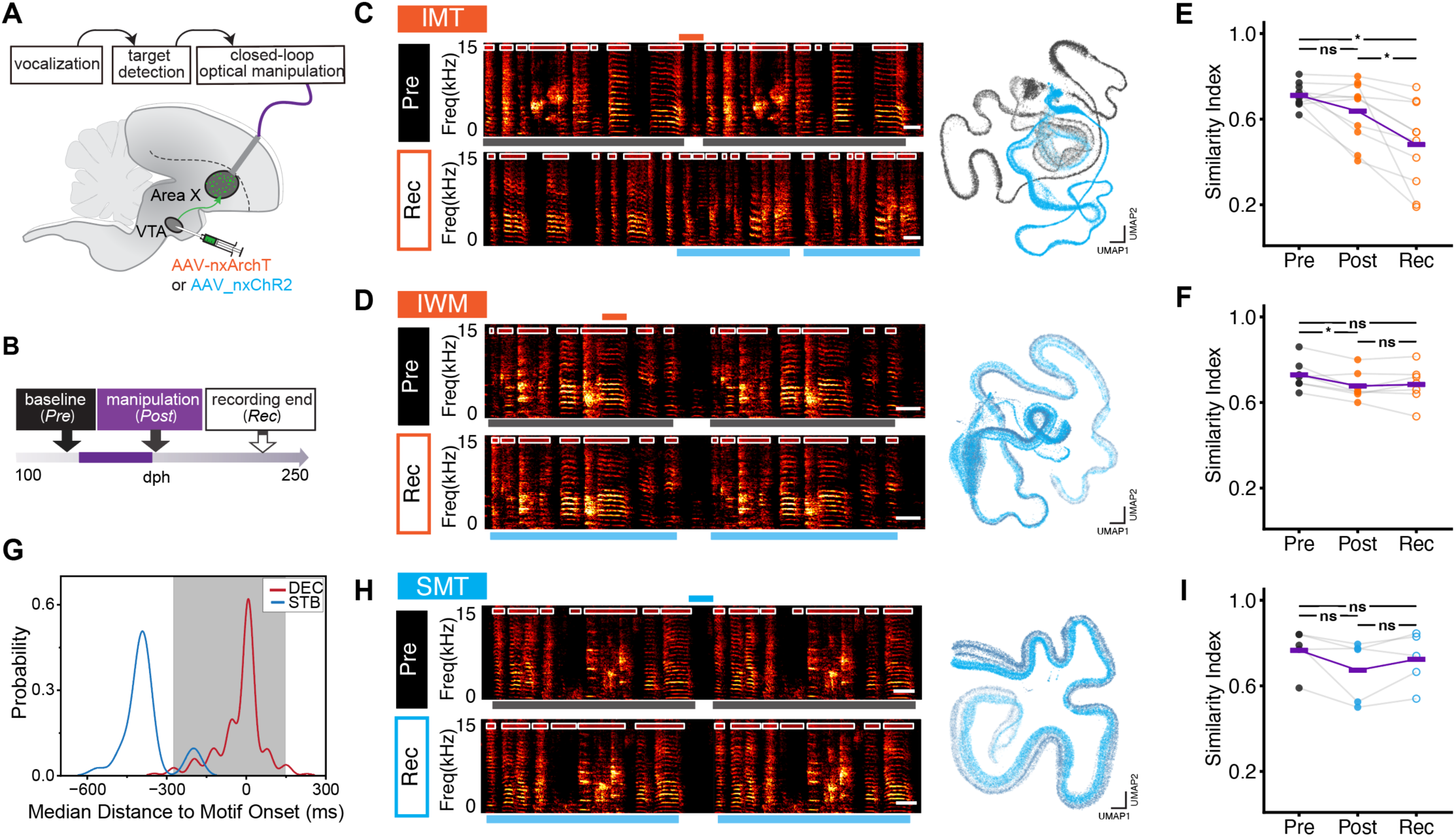
Phasic Inhibition, but Not Stimulation, of Dopamine Signaling at Motif Transitions Causes Song Deterioration. **A)** Experimental design for closed-loop optogenetic manipulation of VTA dopamine terminals in Area X during singing. Birds expressed axon-targeted nxArchT or nxChR2 in VTA “neurons, and optical fibers were implanted over Area X. **B)** Experimental timeline showing baseline song recording (Pre), recording at the end of optogenetic manipulation (Post), and final recording (Rec). **C)** Example inhibition-at-motif-transition (IMT) bird. Left, spectrograms from Pre and Rec song; white boxes mark syllables, orange bar mark inhibition window, and black and blue bars below the spectrograms outline motifs from the Pre and Rec periods, respectively. Right, time-resolved spectrogram embeddings projected into a shared UMAP space show largely non-overlapping Pre and Rec trajectories. Scale bar, 100ms. **D)** Example inhibition-within-motif (IWM) bird. Spectrograms and UMAP trajectories show preserved song structure across Pre and Rec. Scale bar, 100ms. **E)** Similarity index across Pre, Post, and Rec periods for IMT birds. Phasic inhibition at motif transitions caused progressive song deterioration **F)** Similarity index across Pre, Post, and Rec periods for IWM birds. **G)** Bootstrap distributions of inhibition timing relative to motif onset in decrystallized (DEC) and stable (STB) birds. Gray shading indicates the 95% CI for inhibition windows in DEC birds. **H)** Example stimulation-at-motif-transition (SMT) bird. Spectrograms and UMAP trajectories show preserved song structure following optical stimulation; black and blue bars below mark Pre and Rec motifs, respectively, and the blue bar above marks the stimulation window. Scale bar, 100ms. **I)** Similarity index across Pre, Post, and Rec periods for SMT birds. Statistics are summarized in Table 3.

Across 17 adult male zebra finches, we targeted 7,502 ± 1,486 song renditions over 9 ± 1 days. To relate behavioral effects to perturbation timing, we grouped birds according to the empirical timing of optical inhibition relative to each bird’s motif. Inhibition timing was aligned to motif onset and offset and normalized to motif duration. Birds in which the inhibition window fell within the first or last 20% of the motif were classified as inhibition-at-motif-transition (IMT), whereas birds inhibited within the intervening motif body were classified as inhibition-within-motif (IWM).

We compared songs recorded before optogenetic inhibition (Pre, 176 ± 19 dph), on the final day of optical inhibition (Post, 188 ± 21 dph), and at the end of the recovery period (Rec, 196 ± 22 dph; mean ± SEM) (**Fig. 3B**). In an example IMT bird, time-resolved spectrogram embeddings from hundreds of motif renditions in the Pre and Rec periods traced largely non-overlapping trajectories through the same latent acoustic manifold, indicating substantial reorganization of learned vocal structure (**Fig. 3C**). By contrast, the corresponding trajectories in an example IWM bird remained largely overlapping across the same periods (**Fig. 3D**).

Because individual birds varied in the form and magnitude of song change, we quantified behavioral effects using acoustic similarity, sequence similarity, and a combined similarity index (SI) (**fig. S7, A to F; Table 2**). IMT birds showed significant reductions in all three measures by the end of the Rec period relative to Pre, whereas IWM birds showed no significant change in syllable sequencing or combined SI (**Fig. 3, E and F; fig. S8, A to D**). Baseline song similarity did not differ between groups, but by the end of Rec, IMT birds had significantly lower similarity scores than IWM birds (**Table 3**). These results suggested that behavioral deterioration depended on when inhibition occurred relative to motif structure.

**Table 2.**
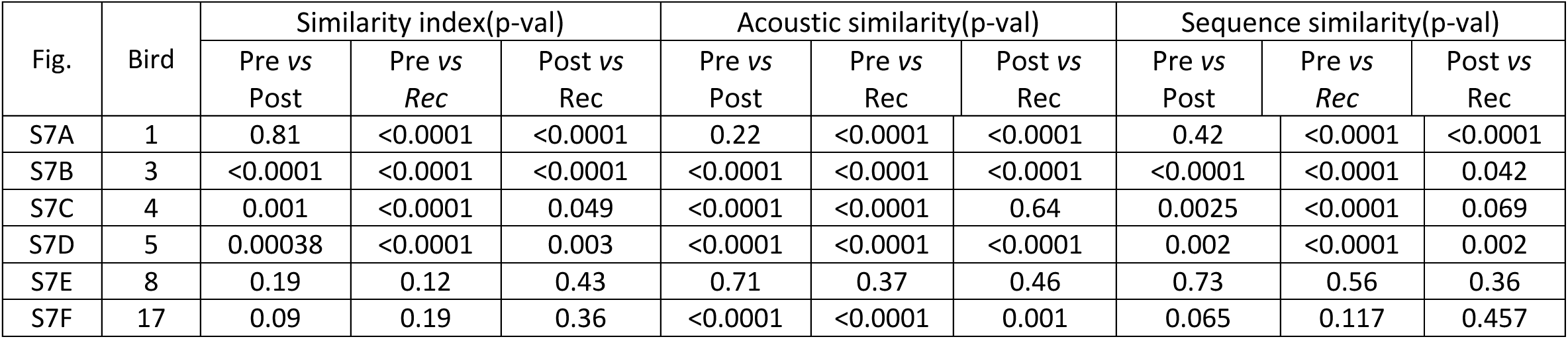
Statistical comparisons of similarity scores across Pre, Post, and Rec periods in ArchT+ birds (fig. S7).

**Table 3.**
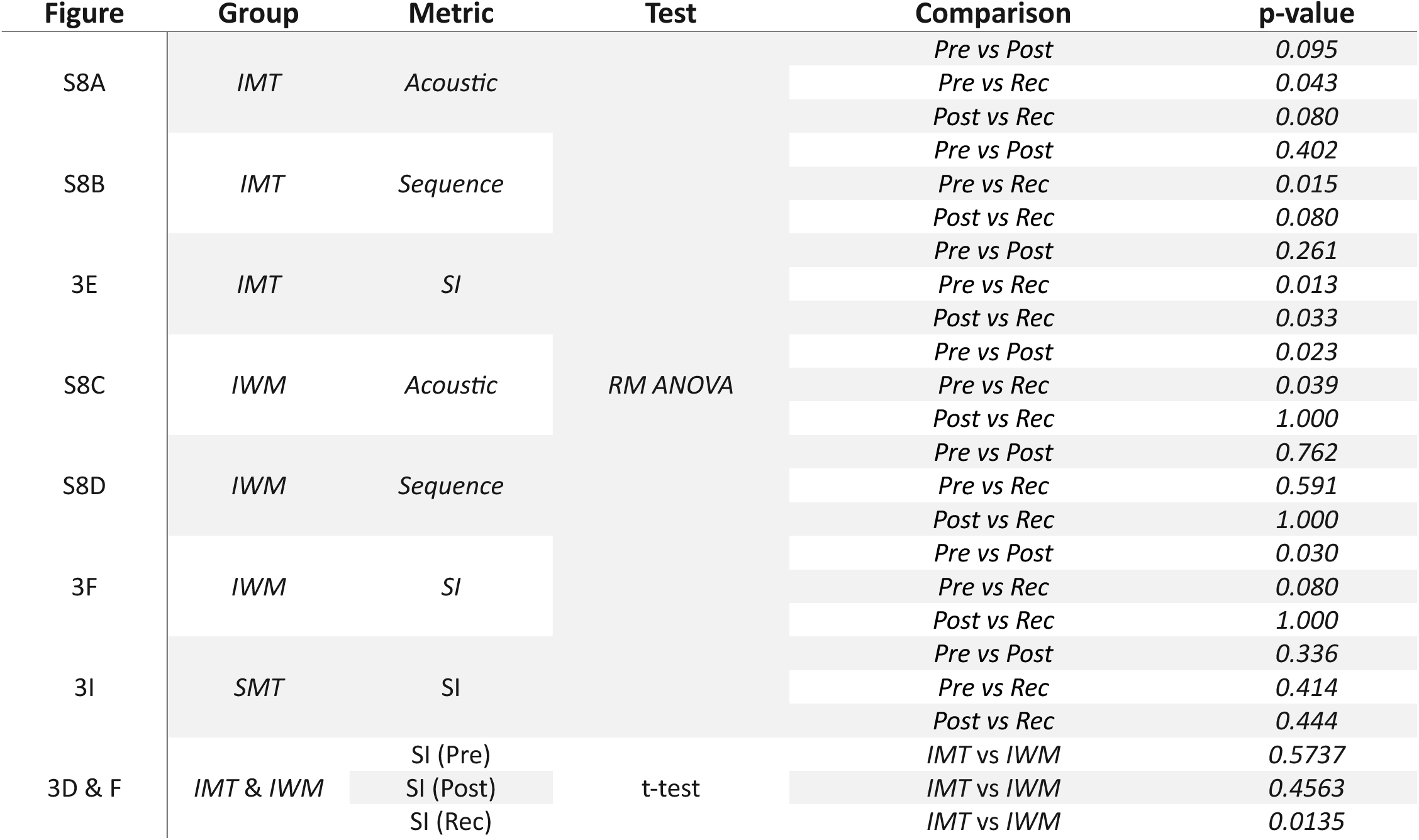
Statistical comparisons for song similarity metrics across Pre, Post, and Rec stages (Fig. 3; fig. S8)

We next tested this relationship at the level of individual birds. By the end of the Rec period, 7 of 10 IMT birds showed significant reductions in SI, whereas only 1 of 7 IWM birds did so, indicating that decrystallization was enriched after motif-transition inhibition (Fisher’s exact test, one-sided; p = 0.036).

Because IMT and IWM birds could differ in variables other than inhibition timing, we next asked which experimental features best predicted endpoint behavioral outcome. To address this question independently of the timing-defined groups, we classified birds by endpoint behavioral outcome as decrystallized (DEC) or stable (STB) based on whether they showed a significant reduction in SI. We quantified illumination timing relative to motif onset and offset, motif duration, daily inhibition rate, total inhibition events, total motif renditions during inhibition days, number of inhibition days, recovery duration, and age at the beginning and end of optical inhibition (**fig. S9, A to J**). Random forest classification identified inhibition timing relative to motif boundaries and recovery duration as the strongest predictors of DEC versus STB outcome (**fig. S9K**). Mutual information analysis further showed that inhibition timing relative to motif onset was the only variable that carried significant information about behavioral deterioration (**fig. S9L**). The combination of motif-onset timing and recovery duration yielded the highest mutual information (**fig. S9M**), consistent with the delayed progression of decrystallization.

Because inhibition timing relative to motif onset emerged as the strongest predictor of behavioral outcome, we next asked whether the behavioral data could localize the vulnerable time window more precisely. Bootstrap analysis of inhibition timing in DEC birds identified a narrow peak in probability centered near motif onset, whereas the distribution in STB birds was shifted away from this window toward times farther from motif boundaries (**Fig. 3G**). This behaviorally inferred vulnerable period aligned with the transient motif-onset dopamine peak observed in adult photometry recordings. Thus, decrystallization was not a general consequence of suppressing dopamine during singing but was specifically associated with perturbation of the motif-onset window in which phasic striatal dopamine signaling is normally strongest. These results indicate that transient dopamine signaling at learned sequence boundaries is required for the long-term stability of crystallized adult song.

### Motif-Onset Dopamine Inhibition Causes Progressive Loss of Learned Song Structure

This motif-onset dopamine transient differed functionally from the within-motif dopamine fluctuations previously linked to ongoing vocal features(*17–23*). In prior work, temporally targeted inhibition or stimulation of these within-motif signals drove rapid, feature-specific adaptive changes in song, with opposite effects depending on excitation or inhibition of dopamine release (*18, 19*). We therefore asked whether increasing dopamine signaling at motif transitions would also alter adult song structure. To test this, we expressed axon-targeted channelrhodopsin (nxChR2) in VTA dopamine neurons and optically stimulated their terminals in Area X at motif transitions using the same temporal protocol as for inhibition (**Fig. 3A**)(*19, 35*). Across five adult birds, stimulation failed to produce detectable song decrystallization or any consistent change in SI (**Fig. 3, H and I**). Thus, unlike bidirectional perturbation of within-motif dopamine fluctuations, increasing dopamine signaling at the motif-transition window did not phenocopy the delayed decrystallization caused by inhibition, suggesting that the motif-onset signal is required for stability but is not sufficient to instruct song restructuring when artificially elevated.

A second distinguishing feature of motif-transition dopamine inhibition was the delayed and progressive nature of the phenotype. Perturbing within-motif dopamine signals associated with ongoing vocal performance produces its largest behavioral effects during the perturbation period, followed by recovery after perturbation ends(*19, 35*). By contrast, song decrystallization after inhibition at the motif-transition window emerged gradually and was often most pronounced during recovery (**Fig. 3, D and F; Table 4**). Thus, the phenotype was not an immediate online consequence of dopamine suppression but developed over days after repeated perturbation of the motif-transition signal.

**Table 4.**
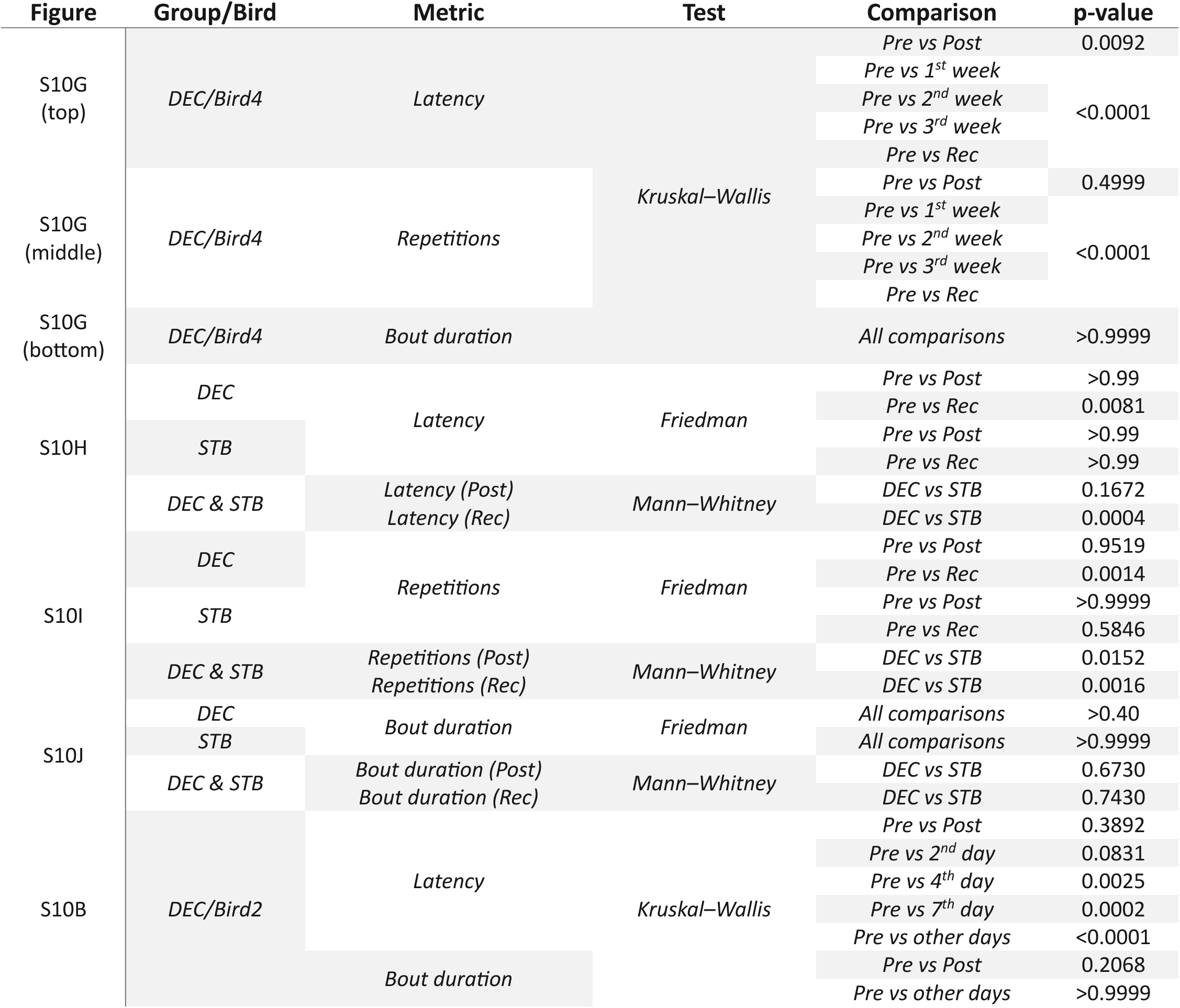
Statistical comparisons for song initiation metrics across Pre, Post, and Rec periods in DEC and STB birds (fig. S10)

In the most strongly affected birds, delayed deterioration included progressive loss of learned vocal elements. Four of eight DEC birds gradually dropped syllables from their song, with increasing omission of syllables from motif ends or replacement of the remaining sequence as recovery progressed (**fig. S10, A and D; fig. S11**). In some birds, nearly all developmentally learned syllables were lost by the end of recovery. These observations indicate that transient dopamine signaling at motif transitions is required to preserve both the acoustic structure of adult song and the learned sequence of its constituent syllables over time.

Previous studies have linked pre-movement dopamine activity to action initiation and movement vigor(*13, 14, 36, 37*). Inhibition at the motif-transition window did not produce comparable acute motor disruption during singing. DEC birds showed no changes in singing rate or song bout duration across the Pre, Post, and Rec periods (**fig. S10, B, G and J**), suggesting that overall song output was preserved. However, all DEC birds progressively increased the number of introductory notes before the first motif in a bout, thereby prolonging the latency to motif initiation (**fig. S10, A, B, and D to J**). This effect was not prominent during optical inhibition itself, but emerged after inhibition ended and grew during recovery, ultimately exceeding that observed in STB birds (**Table 4**). Thus, perturbing the motif-onset dopamine window did not reduce the drive to sing, but disrupted the transition into learned motif production, consistent with a role in maintaining adult vocal sequence organization rather than acutely controlling movement vigor.

## Discussion

Our findings identify a dopamine signal in the sBG that operates at the level of learned sequence boundaries. In contrast to dopamine signals previously linked to syllable-level performance evaluation, motif-onset dopamine appears to support maintenance of a higher-order motor unit: the learned motif(*38*). The temporal specificity of the perturbation suggests that motif transitions are privileged control points in learned song sequences, consistent with proposed basal ganglia roles in action chunking and sequence initiation(*1, 4, 7, 13*). More broadly, adult song decrystallization suggests that sequence stability is not passively retained after learning but instead depends on ongoing dopaminergic signaling at motif onset. In this view, inhibiting motif-onset dopamine in adulthood progressively weakens the integrity of the learned sequence rather than causing only an acute performance disruption. Such destabilization can manifest as loss of previously acquired vocal elements, altered sequence organization, and delayed motif initiation.

The developmental shift of dopamine signaling toward motif onset is consistent with TD-like reinforcement processes operating during natural skill learning, even in the absence of explicit external rewards. In birdsong, internally generated evaluative signals, such as successful motif completion or sensory feedback related to vocal performance, could allow dopamine activity to become progressively anchored to earlier predictive points in the sequence. Its developmental emergence further argues against a purely generic movement-onset response and instead suggests that dopamine signaling becomes organized around motif initiation as learning organizes song into a stable learned sequence(*39, 40*).

Although reinforcement-learning models can account for how an onset-locked predictive signal might emerge, they do not address whether such a signal remains necessary for skilled sequence execution. The delayed decrystallization caused by inhibition at motif transitions suggests that motif-onset dopamine is not simply an online motor trigger but may help preserve the learned organization of the motif across repeated renditions. In a TD-like framework, motif completion could be treated as an internally evaluated outcome whose predictive value becomes anchored to motif onset during learning. Repeated suppression of this boundary-linked signal in adulthood could then disrupt stabilization of the sequence across renditions rather than produce only an immediate performance deficit. The absence of decrystallization after dopamine stimulation at the same window further suggests that the phenotype depends on the learned temporal structure of endogenous dopamine signaling, rather than on perturbation at motif onset per se.

These results broaden the role of basal ganglia circuits after learning. The sBG pathway is often viewed as most important during vocal learning and adaptive plasticity, whereas adult song structure is commonly attributed to premotor cortical circuits that generate the crystallized motif. Our data indicate that basal ganglia dopamine remains functionally engaged after crystallization and contributes to preserving the sequential organization of adult song. This interpretation does not exclude contributions from premotor circuits, nor does it establish that the observed dopamine dynamics implement a canonical TD algorithm. It does, however, support a broader principle that skilled behaviors may require persistent reinforcement-like signals at sequence boundaries to remain stable long after developmental learning is complete.

## Supplementary Figures

**Figure S1.**
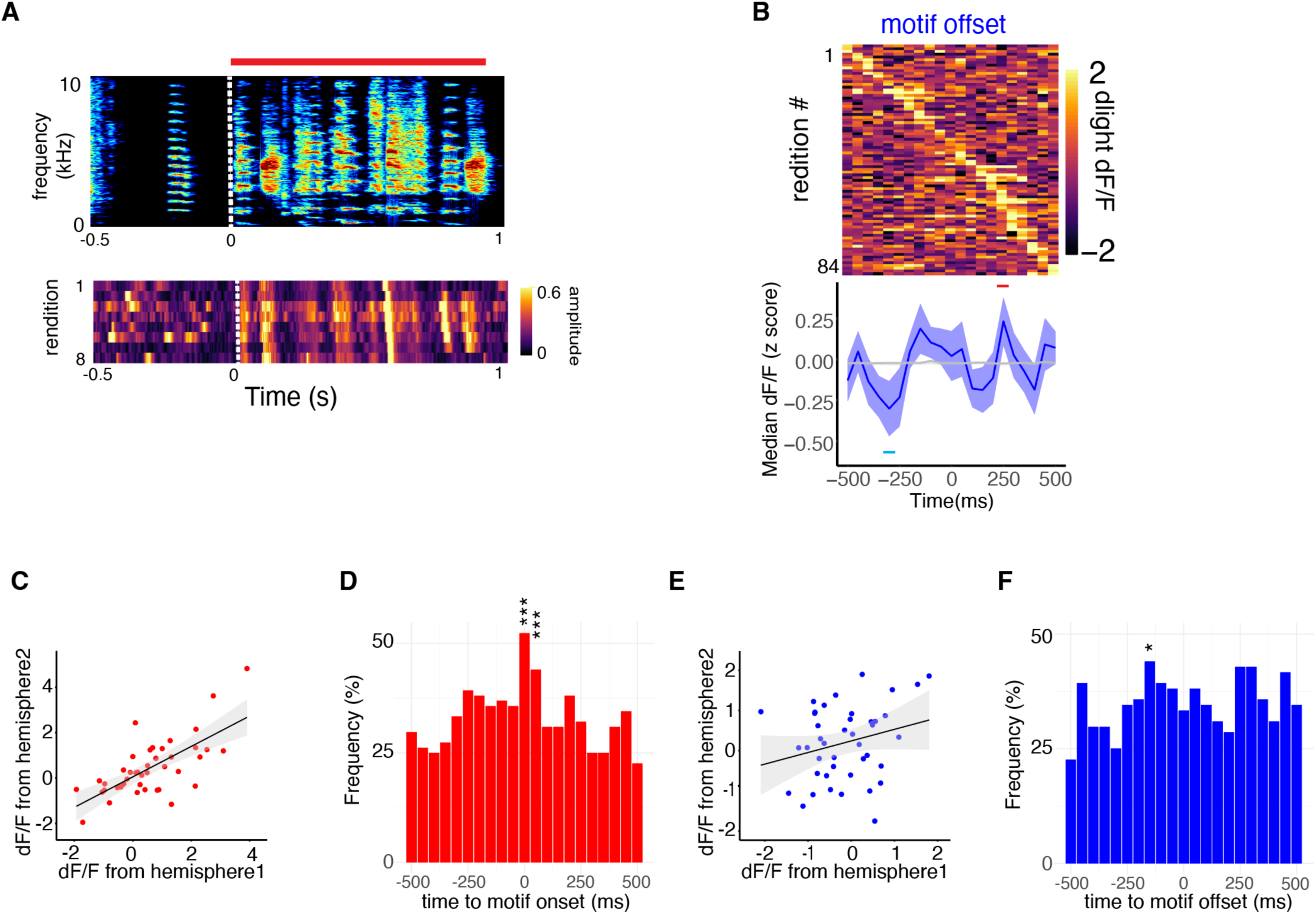
Motif-Aligned Dopamine Dynamics During Directed Singing. **A)** Top, example spectrogram of a song motif aligned to motif onset. The red bar marks the motif, and the white dashed line indicates motif onset. Bottom, sound-amplitude stack plot of eight motif renditions aligned to motif onset, illustrating the temporal consistency of motif structure across renditions. **B)** Trial-by-trial dLight fluorescence aligned to motif offset during directed singing and sorted by peak timing (n = 84 renditions). Bottom, median motif-offset-aligned dLight activity. At motif offset, shuffle analysis detected a significant post-offset peak at 250 ms (red bar; p = 0.010; other time points, p > 0.05) and a significant pre-offset dip at –300 ms (blue bar; p = 0.014). Shaded error indicates ±s.e.m.; grey shading indicates the 95% shuffle CI. **C)** Across-hemisphere correlation of motif-onset-aligned dLight fluorescence. Dopamine signals were significantly correlated between hemispheres near motif onset (R = 0.60, p = 3.7 × 10⁻⁵). **D)** Frequency histogram of detected dLight transients aligned to motif onset. Transients occurred significantly more often near motif onset, with enrichment at 0 ms (p < 0.001) and 50 ms (p = 0.006; shuffle test). **E)** Across-hemisphere correlation of motif-offset-aligned dLight fluorescence. Correlation between hemispheres at motif offset was weak and not significant (R = 0.21, p = 0.19). **F)** Frequency histogram of detected dLight transients aligned to motif offset. No consistent increase in transient frequency was observed around motif offset, although a weak enrichment occurred before offset at –150 ms (p = 0.022; shuffle test).

**Figure S2.**
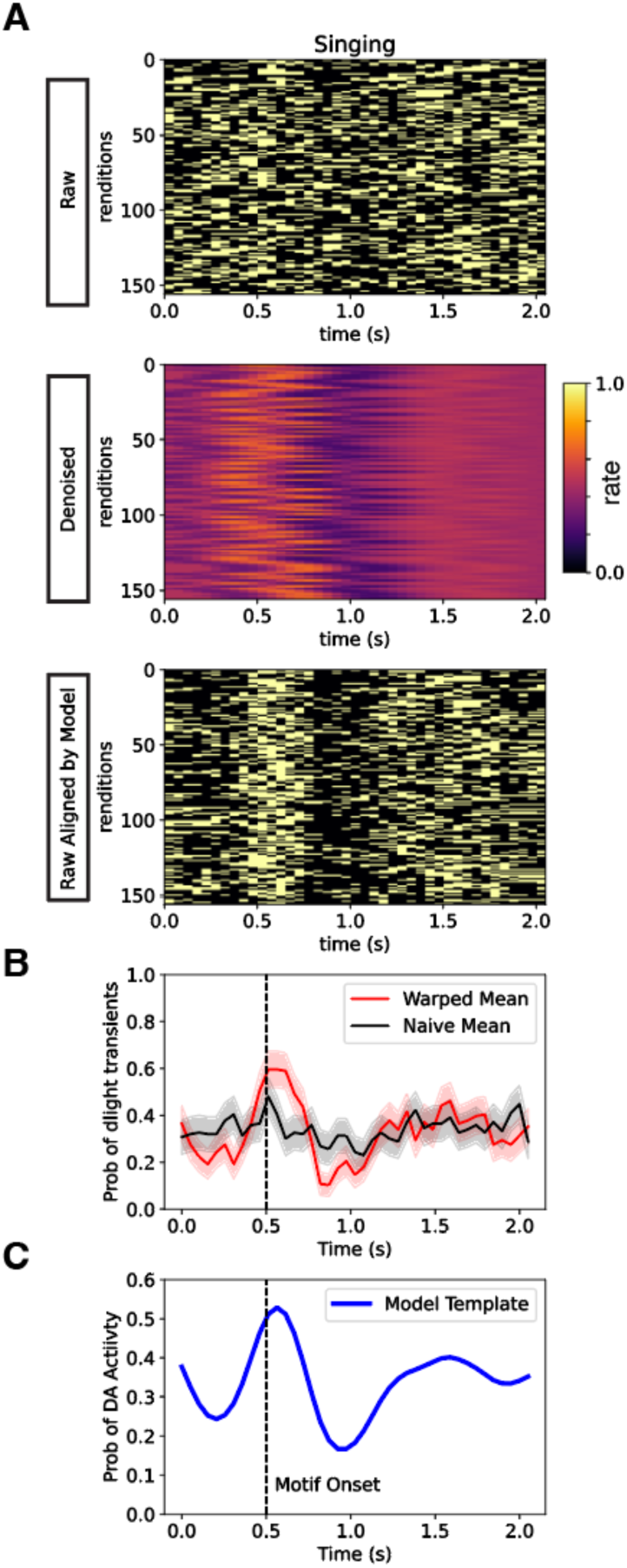
Motif-Aligned Dopamine Transients During Singing. **A)** Trial-by-trial binary dopamine-transient events during singing, shown as raw event rasters (top), denoised transient-rate representations (middle), and raw event rasters realigned by the shift-only time-warping model (bottom). Each row represents one motif rendition. **B)** Mean probability of dopamine transients across motif renditions using naive alignment (black) or shift-only model alignment (red). Shaded areas indicate 95% CI. Dashed vertical line marks motif onset. **C)** Model-derived transient-probability template during singing. Dopamine transients showed a consolidated peak near motif onset followed by a dip within the motif. Dashed vertical line marks motif onset.

**Figure S3.**
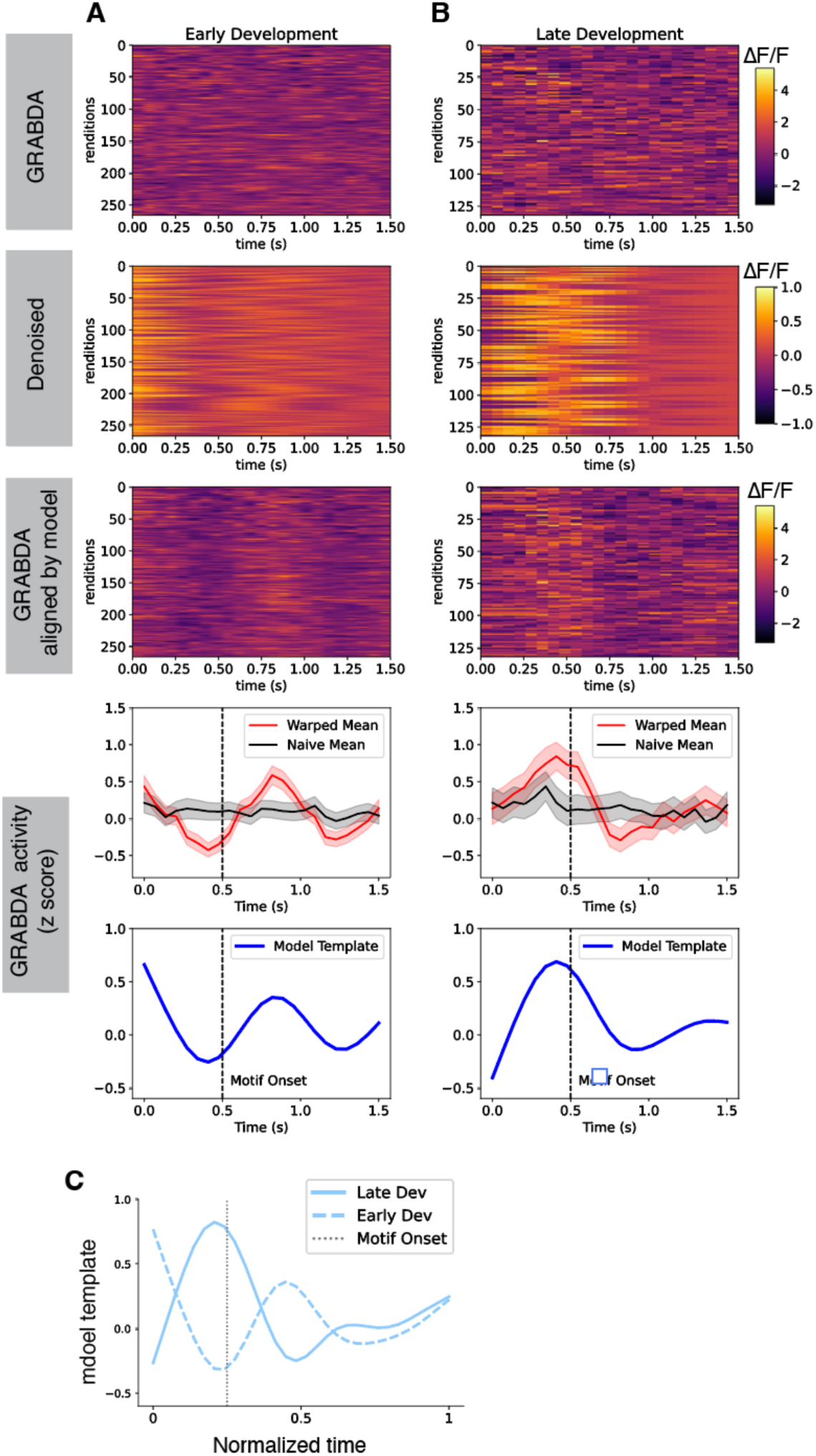
Alignment of GRAB-DA Signals During Song Development. **A)** Trial-by-trial GRAB-DA fluorescence from early-development song, shown as raw fluorescence traces (top), denoised representations (second row), raw traces realigned by the shift-only time-warping model (third row), naive versus model-aligned mean activity (fourth row), and the model-derived activity template (bottom). Each row represents one motif rendition. Dashed vertical line marks motif onset. **B)** Trial-by-trial GRAB-DA fluorescence from late-development song, shown as raw fluorescence traces, denoised representations, model-aligned raw traces, naive versus model-aligned mean activity, and the model-derived activity template as in (**A**). Late-development song showed a consolidated dopamine peak near motif onset. **C)** Model-derived GRAB-DA templates plotted in normalized motif time for early- and late-development song. The dopamine activity peak was offset-biased in early development and shifted toward motif onset by late development.

**Figure S4.**
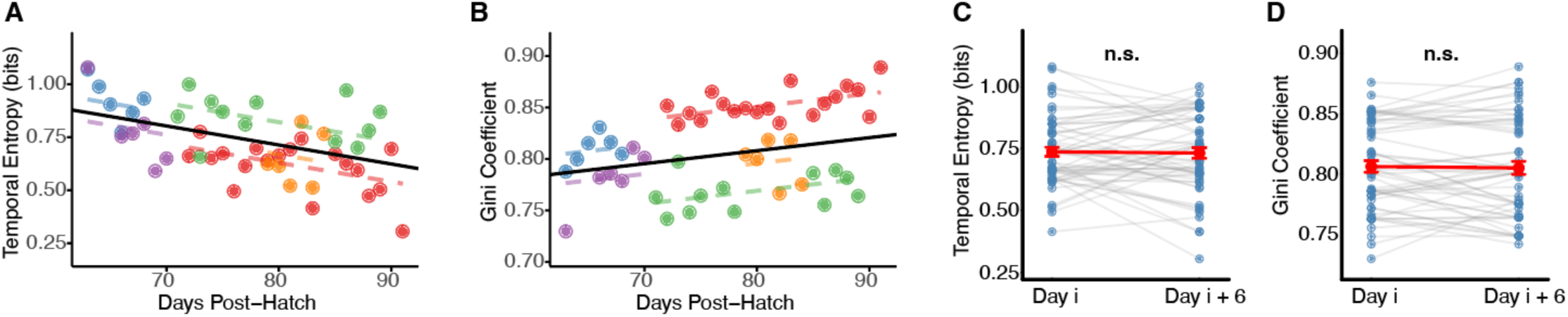
Developmental Changes in Temporal Concentration of Dopamine Transients. **A)** – **B**) Population-level analysis of dopamine-transient temporal concentration across vocal development. Linear mixed-effects models showed that temporal entropy decreased with age (**A**), whereas the Gini coefficient increased with age (**B**) (n = 49 observations from 5 birds, 63 to 91 dph). Each colored dot represents the mean value for one bird at a given age; colors indicate individual birds. Black lines show population-level fixed effects from LMMs with bird identity as a random intercept. **C)** – **D**) Paired comparison of temporal concentration metrics between the first day (Day i) and last day (Day i + 6) of 6-day sliding windows. Gray lines connect paired observations from individual birds within the same temporal window (n = 57 paired observations from 5 birds across 24 windows). Red symbols indicate group means ± SEM. Temporal entropy (**C**) and Gini coefficient (**D**) did not change significantly within 6-day windows. Statistics are summarized in Table 1.

**Figure S5.**
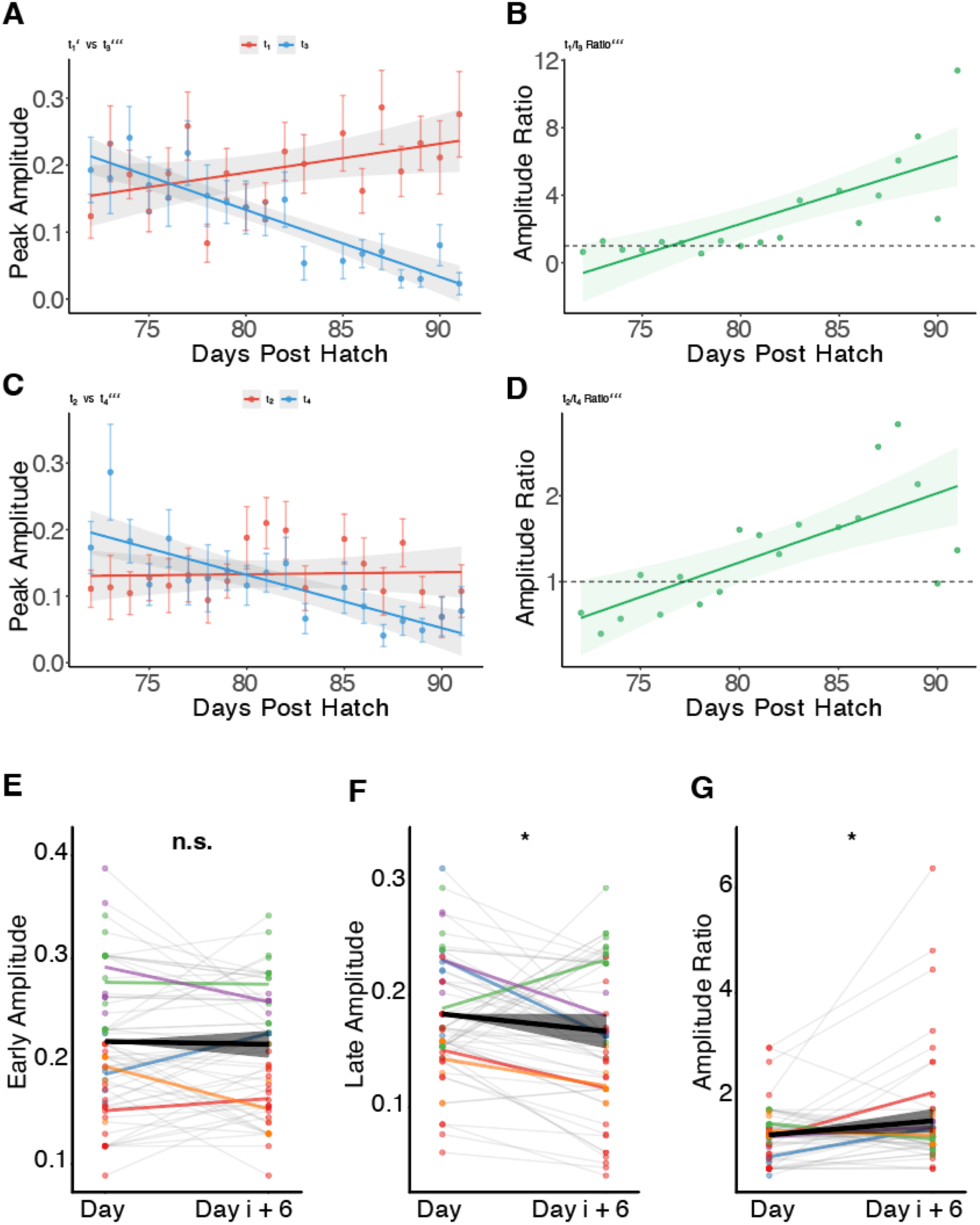
Dopamine Amplitude Shifts Across Motif Segments During Vocal Development. **A)** – **D**) Representative example from a single bird showing amplitude redistribution across different pairs of motif segment. (**A**) Peak amplitudes for early (t_1_, red) and late (t_3_, blue) segments across development. Early amplitude increased significantly while late amplitude decreased significantly. (**B**) Early/late amplitude ratio (t_1_/t_3_) increased significantly across development. (**C**) Peak amplitudes for early (t_2_, red) and late (t_4_, blue) segments showing developmental trajectory. Early amplitude showed no significant change while late amplitude decreased significantly. (**D**) Early/late amplitude ratio (t_2_/t_4_) increased significantly across development. **E)** – **G**) Population-level within-window paired analysis using mixed-effects models across 6-day sliding windows. Points: individual observations colored by animal. Gray lines: individual observations within each window (animal × window combinations). Colored lines: mean trajectory for each animal across all windows. Black line with ribbon: population-level trend with 95% CI from fixed effects. (**E**) Early amplitude showed no significant within-window change. (**F**) Late amplitude decreased significantly within windows. (**G**) Amplitude ratio increased significantly within windows. Statistics are summarized in Table 1.

**Figure S6.**
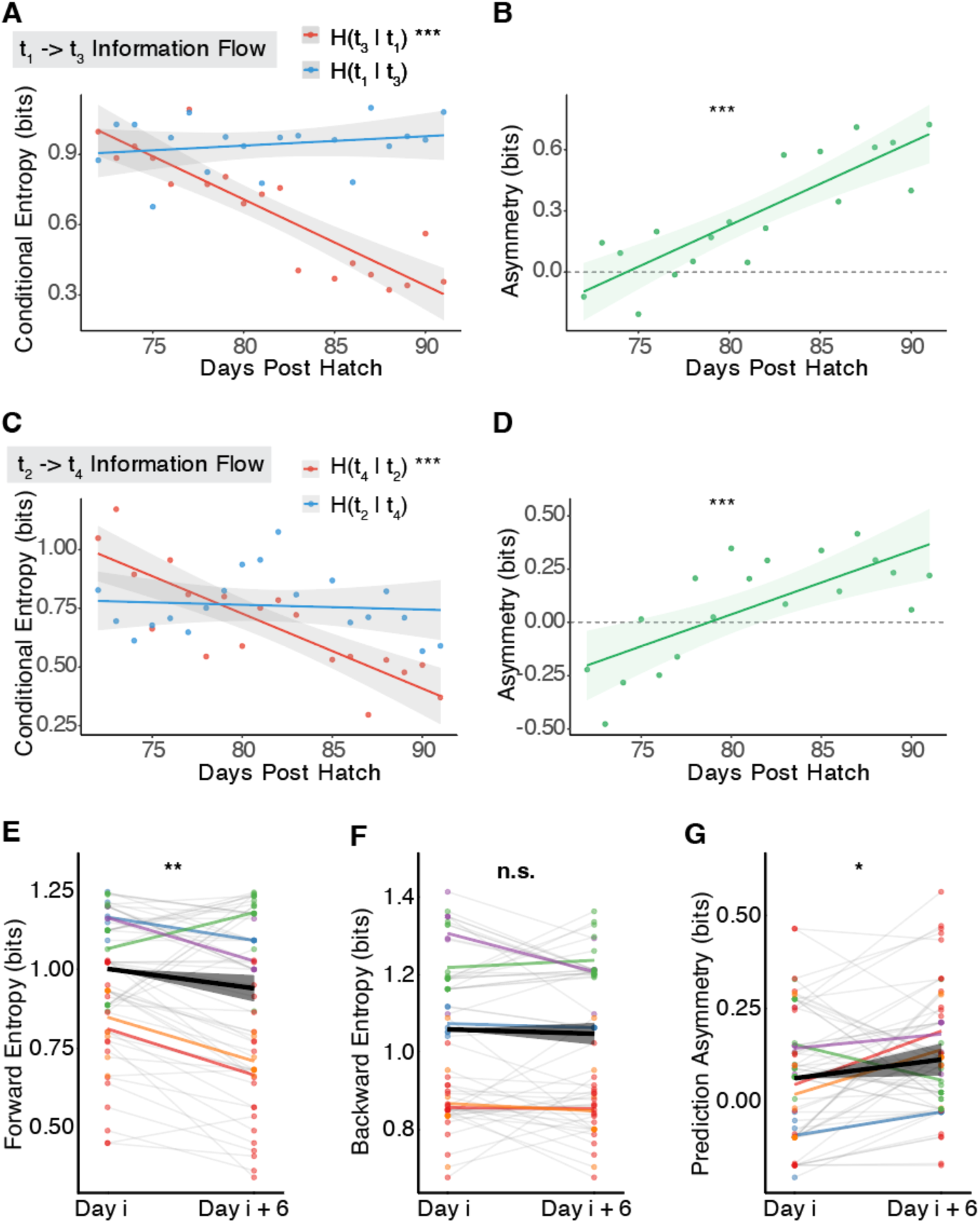
Directional Information Flow Pattern across Different Motif Segments During Vocal Development. **A)** – **D**) Representative example from a single bird showing directional information flow between different pairs of motif segments. (**A**) Conditional entropies for forward (H(t_3_|t_1_), red) and backward (H(t_1_|t_3_), blue) prediction across development. Forward entropy decreased significantly, indicating improved predictability of t_3_ from t_1_, while backward entropy showed no significant change. (**B**) Prediction asymmetry (H(t_1_|t_3_) - H(t_3_|t_1_)) increased significantly across development, with dominance crossover at DPH 74.4. (**C**) Conditional entropies for forward (H(t_4_|t_2_), red) and backward (H(t_2_|t_4_), blue) prediction showing developmental trajectory. Forward entropy decreased significantly, while backward entropy showed no significant change. (**D**) Prediction asymmetry (H(t_2_|t_4_) - H(t_4_|t_2_)) increased significantly across development, with dominance crossover at DPH 78.7. **E)** – **G**) Population-level within-window paired analysis using mixed-effects models across 6-day sliding windows. Points: individual observations colored by animal. Gray lines: individual observations within each window (animal × window combinations). Colored lines: mean trajectory for each animal across all windows. Black line with ribbon: population-level trend with 95% CI from fixed effects. (**E**) Forward entropy decreased significantly within windows, indicating improved forward predictability. (**F**) Backward entropy showed no significant within-window change. (**G**) Prediction asymmetry increased significantly within windows, demonstrating a shift toward forward prediction dominance. Statistics are summarized in Table 1.

**Figure S7.**
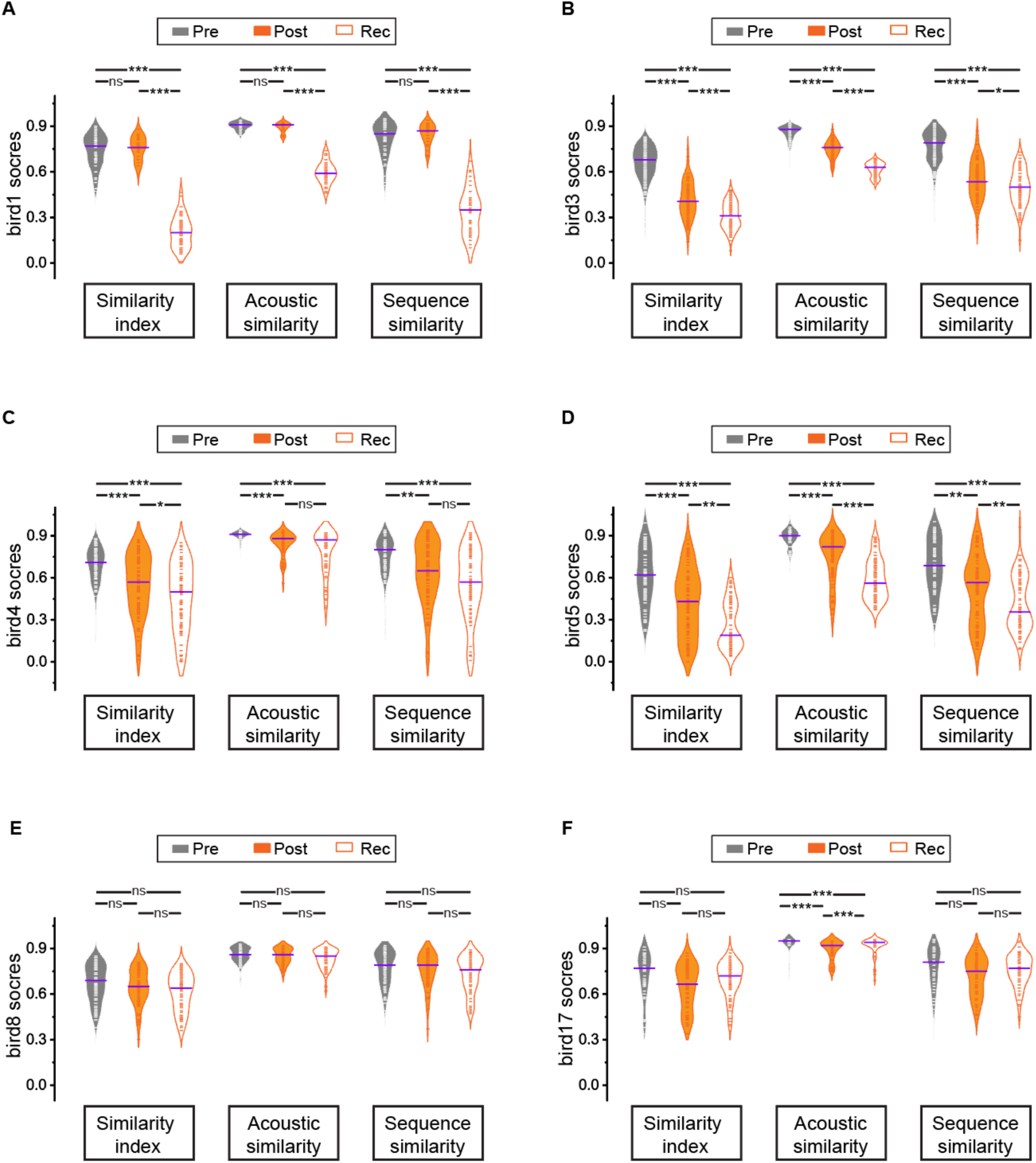
Individual Variability in Similarity Metrics across Pre, Post, and Rec Periods in ArchT+ Birds. **A)** – **F**) Violin plots of similarity index, acoustic similarity, and sequence similarity scores for individual ArchT+ birds (Bird ID: 1, 3, 4, 5, 8, 17) recorded at Pre, Post, and Rec periods. Lines indicate median scores. Statistical comparisons for each bird are shown in Table 2(Mood’s tests).

**Figure S8.**
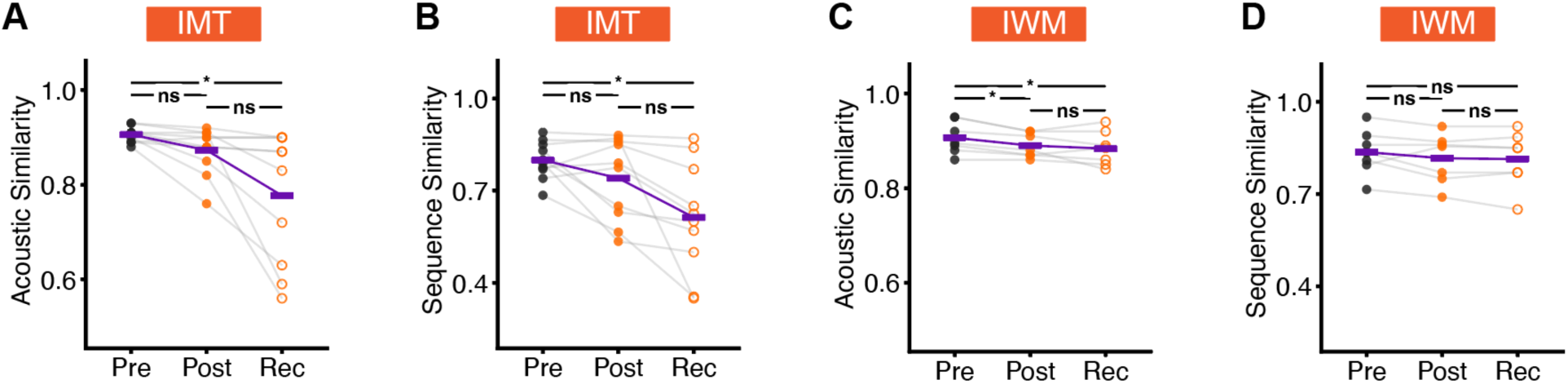
Acoustic and Sequence Similarity After Dopamine Inhibition at Motif Transitions or Within Motifs. **A)** – **B**). Acoustic similarity (**A**) and sequence similarity (**B**) across Pre, Post, and Rec periods in IMT birds. IMT birds showed progressive reductions in both acoustic and sequence similarity after optogenetic inhibition. **C)** – **D**). Acoustic similarity (**C**) and sequence similarity (**D**) across Pre, Post, and Rec periods in IWM birds. IWM birds showed limited disruption, with no progressive decline in sequence similarity. Gray lines indicate individual birds; purple lines indicate group means. Statistics are summarized in Table 3.

**Figure S9.**
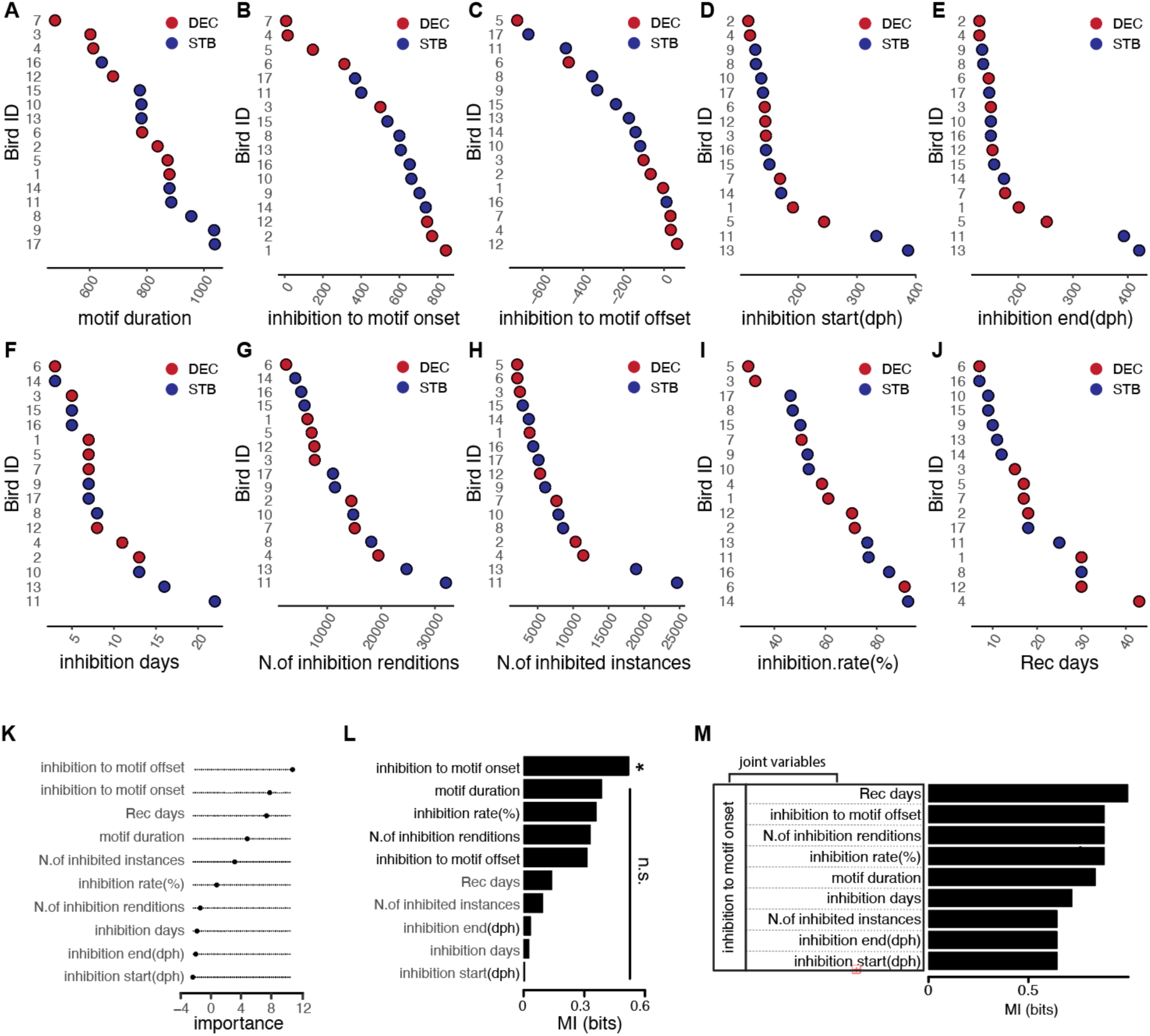
Experimental Variables Associated with Song Decrystallization after Dopamine Inhibition. **A)** – **J**) ArchT+ birds sorted by experimental and behavioral variables and labeled by endpoint behavioral outcome. Red points indicate decrystallized (**DEC**) birds; blue points indicate stable (**STB**) birds. Variables include motif duration (**A**), inhibition timing relative to motif onset (**B**), inhibition timing relative to motif offset (**C**), inhibition start age (**D**), inhibition end age (**E)**, number of inhibition days (**F**), number of inhibited song renditions (**G**), number of inhibited instances (**H**), inhibition rate (**I**), and recovery duration (**J**). **K)** Random forest feature importance for classifying **DEC** versus **STB** birds. Inhibition timing relative to motif offset and onset, and recovery duration ranked among the strongest predictors. Points indicate feature-importance estimates. **L)** Mutual information (**MI**) between individual experimental variables and **DEC**/**STB** outcome. Inhibition timing relative to motif onset was the only variable significantly associated with behavioral outcome (p = 0.032; all other variables, p > 0.05). **M)** Joint-variable **MI** analysis using inhibition timing relative to motif onset paired with each additional variable. The largest MI was observed for motif-onset timing combined with recovery duration, consistent with progressive decrystallization after perturbation near motif onset.

**Figure S10.**
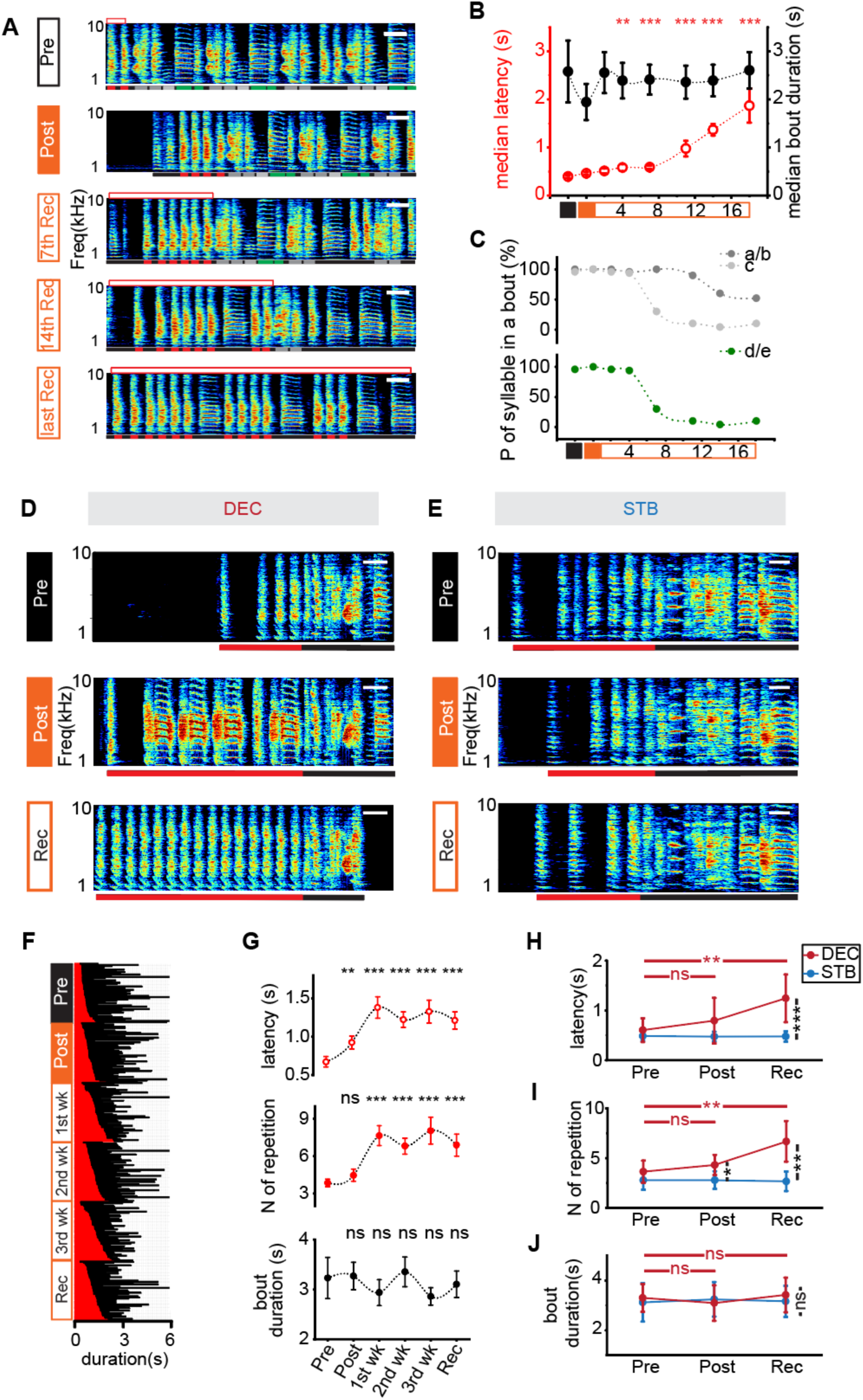
Progressive Song Changes Following Phasic Inhibition of Dopamine Signaling at Motif Transitions. **A)** Representative spectrograms from a **DEC** bird across Pre, Post, 7th Rec, 14th Rec, and last Rec sessions. Red boxes above each spectrogram indicate the latency to motif initiation; Red bars mark introductory notes; green bars mark syllables that were progressively lost; gray bars mark retained song elements. Scale bar, 100 ms. **B)** Quantification for the **DEC** bird shown in (**A**). Motif-initiation latency increased progressively during recovery, whereas bout duration remained stable. Red, latency to motif initiation; black, bout duration. **C)** Probability of syllable occurrence for the **DEC** bird shown in (**A**). Specific learned syllables were progressively omitted during recovery. **D)** Example spectrograms from a **DEC** bird across Pre, Post, and Rec periods. Red bars mark introductory notes; black bars mark the first motif. Scale bar, 100 ms. **E)** Example spectrograms from a **STB** bird across Pre, Post, and Rec periods. Red and black bars mark introductory notes and the first motif, respectively. Scale bar, 100 ms. **F)** Example song bouts from the **DEC** bird shown in (**D**), illustrating progressive prolongation of motif-initiation latency across recovery. Red marks latency to motif initiation; black marks bout duration. **G)** Quantification for the **DEC** bird shown in (**D**). Motif-initiation latency and introductory-note repetitions increased after inhibition and during recovery, whereas bout duration remained unchanged. **H)** – **J**). Group analyses of motif-initiation latency (**H**), number of introductory-note repetitions before the first motif (**I**), and bout duration (**J**) across Pre, Post, and Rec periods in **DEC** and **STB** birds. **DEC** birds showed progressive increases in latency and introductory-note repetitions, whereas bout duration remained unchanged. **STB** birds showed no comparable changes. Data are mean ± SEM. Statistics are summarized in Table 4.

**Figure S11.**
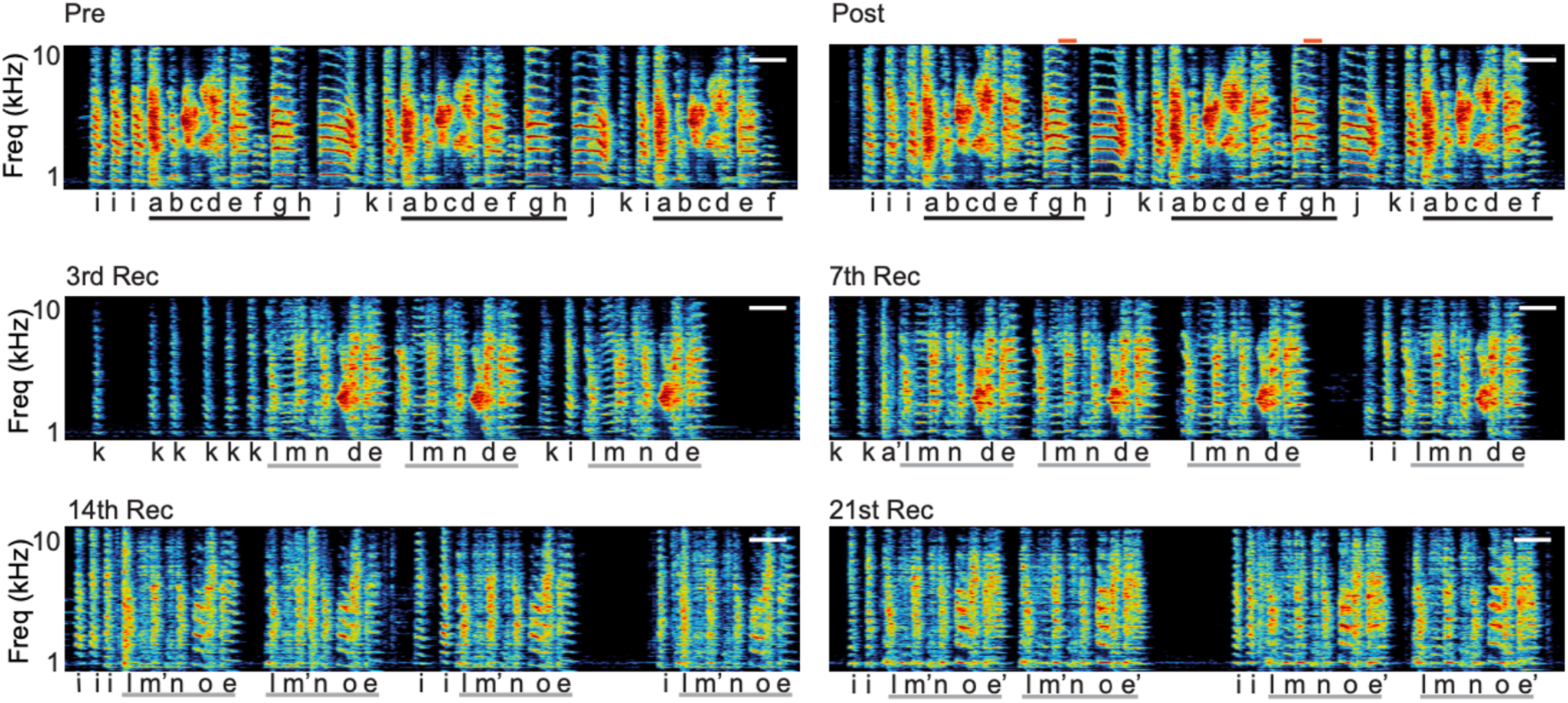
Progressive Loss of Learned Song Structure After Motif-Transition Dopamine Inhibition. Expanded example from the **DEC** bird shown in Fig. 3C, with additional recovery sessions and syllable annotations. Spectrograms show progressive omission or replacement of learned vocal components from Pre and Post through 3rd, 7th, 14th, and 21st Rec sessions. Individual syllables are annotated alphabetically; i marks introductory notes, and syllable variants are indicated by primes (aʹ, eʹ, mʹ). Black lines denote baseline motifs, and gray lines indicate altered motifs. Scale bars, 100 ms.

## METHODS

### Animals

Experiments were performed on 27 adult male zebra finches (*Taeniopygia guttata*) raised in our breeding facility and housed with their parents until at least 50 days of age. During experiments, birds were housed individually in sound-attenuating recording chambers (Med associates) on a 12/12 h day/night cycle and were given ad libitum access to food and water. All procedures were performed in accordance with established protocols approved by the UT Southwestern Medical Center Animal Care and Use Committee.

### Fiber photometry

#### Data sources

Fiber photometry data included both an adult dataset from our original experiments and an open-source juvenile dataset. To acquire the adult data, dopamine sensor dLight1.3b was delivered to Area X of adult male birds via AAV-CAG-dLight1.3b injection (Addgene, #125560). Bilateral implantation of fiber optic cannulae (Thorlab, CFML14U) was performed in the dorsal Area X, and recordings were conducted eight weeks post-injection (240 ± 42 dph). The dLight fluorescence was recorded using fiber photometry (FP3002, Neurophotometrics) controlled via Bonsai™ while male birds were singing alone or toward a female bird. The photometry system used a 470 nm LED for excitation and a 415 nm LED for isosbestic control, with signals sampled at 40 Hz during recording sessions. Virus expression and fiber placement were verified by immunostaining against GFP at the experiment’s end. The juvenile dataset was obtained from an open-source resource ([Duke University Library Research Data Repository](https://research.repository.duke.edu/record/319)), featuring juvenile zebra finch (63-91 days post-hatch) with AAV2/9-GRAB-DA2m sensors injected into the Area X between 33-38 days post-hatch.

#### Signal processing pipeline

Raw photometry data were analyzed using customized functions (VNS, v0.0.0.9, available at GitHub). Datasets underwent baseline correction using an adaptive iteratively reweighted penalized least squares (airPLS) algorithm and ΔF/F calculation through linear regression alignment of control (415 nm) to signal (470 nm) wavelengths. Normalized dopamine fluorescence signal (ΔF/F) was defined as (470 nm signal – fitted 415 nm signal)/ (fitted 415 nm signal)(*41*). Dopamine transients were defined as high amplitude events with a local maximum greater than two times the median absolute deviation (MAD) above the median of the moving window. The processing approaches differed based on specific datasets and recording characteristics, as detailed below.

#### Session-based processing

Adult photometry recordings were processed using whole-session standardization, where signals were normalized using session-wide mean values and standard deviations. This approach was suitable for the restricted recording duration (≤3 minutes per session). To explore dopamine activity patterns in adult birds, we initially employed a supervised alignment approach. Photometry data were manually segmented based on detected motif timestamps, aligned to motif onset or offset, and averaged across motif renditions. Shuffle tests (n=1000 permutations) were conducted to identify time frames with characteristic activity, comparing observed dLight traces against randomized temporal distributions. To address limitations of supervised alignment, such as potential misalignment with behavioral events and temporal dynamics masking, we implemented an unsupervised shift-only time warping model(*29*). This model was applied to extended time windows covering entire motifs, allowing the characterization of dopamine dynamics across complete behavioral sequences independent of predefined alignment point.

#### Epoch-based processing

Juvenile recordings were processed using epoch-based methods to accommodate longer sessions (5-10 minutes) and varied vocalization contexts. Customized functions were used to extract song bouts from audio recordings, which were aligned on the basis of a preselected syllable template from the first motif in each bout (see Methods, “Longitudinal tracking of song change”). Epoch windows were dynamically calculated using quantile-based selection, capturing ∼90-100% of typical vocalization contexts while minimizing overlap between epochs. Photometry recordings were standardized using epoch-wide mean values and standard deviations. Individual epochs were aligned to anchor time stamps for each animal. Given that initial song motifs were well aligned, subsequent analysis focused on the 600-800 ms windows corresponding to motif boundaries. To assess the amplitude shift and calculate conditional entropy across development, this window was further divided into 3-4 segments of 200 ms each. Only the unsupervised method was used to analyze dopamine activity at individual time points during song development.

### Linear mixed-effects models (LMMs)

To examine Temporal Difference (TD) learning signatures, we assessed changes in the timing and amplitude of dopamine transients within the song motif across development using complementary TD learning metrics. To evaluate their developmental trajectories, we employed linear mixed-effects models to analyze individual metrics for population analysis. This approach accounts for the hierarchical data structure with repeated measurements within individuals across multiple developmental timepoints (days post-hatch, dph).

For all TD learning metrics analyses, we fit models of the general form:

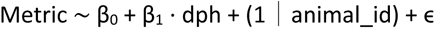

where the outcome variable was the specific TD metric (e.g., median centroid position or backward shift rate). We fitted developmental age (dph) as a fixed effect to capture population-level developmental trajectories, while including a random intercept for individual animals to account for baseline differences between birds. This approach allows for individual variation in baseline values while estimating the common developmental patterns across the population.

Models were fitted using the lme4 package, with degrees of freedom and p-values calculated using the lmerTest package(*42, 43*). For each model, we extracted variance components to quantify between-animal and within-animal (residual) variation, and calculated intraclass correlation coefficients (ICC) to determine the proportion of total variance attributable to individual differences.

To ensure that our developmental LMM results were not driven by uneven sampling across animals, we performed two complementary robustness analyses.

#### Inverse Sample Size Weighting

We refit each LMM using per-observation weights equal to the inverse of the number of observations contributed by each animal (weight = 1/n_obs). This approach equalizes the overall influence of each bird while retaining all available data. Weighted models followed the same structure as the primary unweighted models. Estimates for both weighted and unweighted analyses are reported in Table 1.

#### Bootstrapped Balanced Resampling

To further assess robustness, we generated 1,000 balanced datasets by sampling each animal down to the minimum number of available observations. Each balanced dataset was fit with the same LMM, and the resulting distribution of slope estimates was used to evaluate stability under equal sampling.

Across both approaches, fixed-effect estimates were consistent with the primary unweighted analyses. Because all methods yielded the same developmental conclusions, unweighted LMMs were used for all plots and main-text reporting.

### Temporal shift metrics

#### Peak Timing

We calculated the temporal center of mass (centroid) of the dopamine transients within the motif window for each epoch and then computed the median centroid position for each dph in individual animals. Centroid is calculated as:

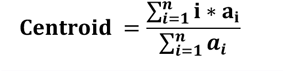

#### Backward Shift Rate

For each dph, we calculated the backward shift rate as the percentage of dopamine activity shift from the motif offset toward onset:*R* = (*n* − *p*)/(*n* − 1) × 100%, where *n* represents the total number of time bins per epoch, and *p* is calculated as the average centroid for individual dph. This metric quantifies the progression of dopamine activity from later to earlier time points, with 0% indicating peak activity at the motif end and 100% indicating peak activity at the motif onset.

For population analysis, we modeled median centroid position (converted to milliseconds from motif onset) or backward shift rate (percentage) as the dependent variable, with dph as a fixed effect and animal identity as a random effect. A significantly negative slope for peak timing (β_1_ <0) and/ or a significantly positive slope for backward shift rate (β_1_ >0) indicated a shift of dopamine peak activity toward earlier time bins with development, consistent with TD learning predictions.

### Temporal sharpening metrics

#### Entropy

For each epoch, we calculated the temporal entropy of dopamine transient activity within the motif:

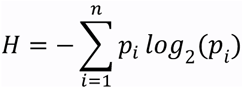

where *p_i_* is the proportion of activity in time bin *i*, and *n* is the total number of time bins per epoch. Lower entropy indicates activity concentrated in fewer time bins (more focused), while higher entropy indicates activity spread across many bins (more dispersed).

#### Gini Coefficient

For each epoch, we calculated the Gini coefficient to quantify inequality in the temporal distribution:

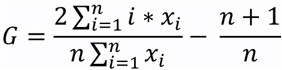

Where *x_i_* represents the sorted activity values, and n is the total number of time bins per epoch. The Gini coefficient ranges from 0 (equal distribution across all bins) to 1 (all activity in one bin). Higher values indicate greater temporal concentration.

For population analysis, we calculated mean entropy and mean Gini coefficient for each developmental age. We then modeled these metrics as dependent variables with dph as a fixed effect and animal identity as a random effect. A significantly negative slope for entropy (β_1_<0) and/or a significantly positive slope for Gini coefficient (β_1_>0) indicated temporal sharpening—dopamine activity becoming more precisely timed with development, consistent with TD learning predictions.

### Amplitude shift metrics

To quantify the redistribution of dopamine peak activity over development, we segmented each song motif into 200-ms intervals and extracted peak amplitudes from each segment using a peak-detection algorithm with a detection threshold defined as a z score exceeding 2 MAD above the ΔF/F median. For each segment, mean peak amplitudes were computed per animal and for each dph. For any pair of motif segments, we further defined a segment amplitude ratio as the mean amplitude of the earlier segment divided by that of the later segment.

For developmental trajectory analyses, we focused on the first and last segments of the motif, denoted as t_1_ ("early") and t_n_ (("late", either t_3_ or t_4_, depending on motif duration). For each animal, we fitted linear regression models to test for changes in early amplitude, late amplitude, and their ratio across dph. At the population level, we used LMMs with dph as a fixed effect and animal identity as a random intercept.

For a more systematic analysis of short-term developmental dynamics, we implemented a sliding window approach that used all available motif segment pairs separated by two segments (e.g., t_1_ vs t_3_, t_2_ vs t_4_), allowing individual birds to contribute multiple early/late comparison pairs within the same song motif. Linear regressions were performed for each pairwise segment comparison within individual animals. For population-level analysis, we used LMMs including motif segment pair and animal identity as random effects and dph as a fixed effect, thereby accounting for repeated measures and potential data imbalance across individuals.

A significantly positive slope(β_1_>0) in early amplitude or the early/late ratio, or a significantly negative slope(β_1_<0) in late amplitude, was interpreted as evidence for a developmental shift toward earlier peak activity, consistent with TD learning mechanisms.

### Directional conditional entropy analysis

To assess changes in predictive information flow across the motif during development, we calculated directional conditional entropy metrics based on dopamine transients used in the temporal analysis above. Each motif was divided into equal-duration segments (as in amplitude analysis), and we focused on the first (t_1_, “early”) and last (t_n_, “late”) segments to quantify information transfer from early to late events and vice versa.

For each animal at each developmental stage, we computed the conditional entropy of the target segment given the source segment—specifically, H (t_n_| t_1_) (forward), H (t_1_| t_n_) (backward), and calculated the asymmetry as their difference (asymmetry = H (t_1_| t_n_) – H (t_n_| t_1_)). These entropy metrics were computed from dopamine transient counts in the respective segments using:

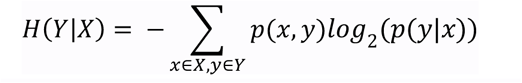

where *p*(*x*, *y*) is the joint probability of observing transient counts *x* in the source segment and *y* in the target segment, and *p*(*y*|*x*) is the conditional probability of *y* given *x*. Lower conditional entropy indicates higher predictability, and increased asymmetry reflects a stronger directional information flow from early to late events.

For developmental analysis in individual animals, we fit linear regression models to test for trends in forward conditional entropy, backward conditional entropy, and asymmetry prediction across dph. At the population level, we applied LMMs to each metric, with dph as a fixed effect and animal identity as a random intercept. A significant negative slope (β_1_ <0) in forward conditional entropy (decreasing unpredictability of late events given early events) or a significant positive slope (β_1_>0) in asymmetry (increasing predictive dominance from early to late), was interpreted as evidence for maturation of predictive coding over development.

### Within-window paired analysis

To detect short-term learning-related changes while controlling for baseline differences between individuals and ensuring repeated assessment across the entire experimental timeframe, we implemented a sliding window approach using the previously described TD metrics. Recording sessions were segmented into overlapping 6-day windows, with each window incremented by 1 day, resulting in 24 windows spanning the experimental period. For each animal and each window, we calculated the metric value on the first and last day of the window, yielding paired observations that reflect changes within consistent time frames. For each metric, we then computed the within-window change across all animals and windows.

For temporal shift metrics, we assessed whether there were systematic short-term developmental trends by applying the Wilcoxon signed-rank test to test the null hypothesis that the median change was zero across all paired observations. This non-parametric approach was chosen due to its robustness to outliers and distributional assumptions.

For amplitude shift and directional conditional entropy metrics, we employed linear mixed-effects models rather than paired tests because one bird contributed two types of time segment comparisons (t_1_ vs t_3_ and t_2_ vs t_4_) while all other birds provided only one pair (t_1_ vs t_3_), resulting in an unbalanced dataset. To ensure statistical power and prevent any single bird from dominating the results, we used the following mixed-effects model:

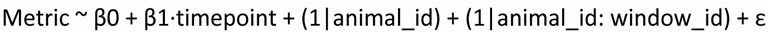

where timepoint is coded as 0 for the first day and 1 for the last day of each window. The model includes random intercepts for each animal to account for baseline differences, and for each window within animals to capture window-specific variation. The fixed effect β1 estimates the average within-window change across all animals and windows, testing whether the mean change is significantly different from zero. This approach properly weights contributions from each animal and provides unbiased population-level estimates despite the data imbalance.

### Optogenetic manipulation

All procedures were performed as reported previously(*19*). Briefly, male birds were randomly assigned bilateral injection of AAV-nxArchT or -nxChR2 in VTA at ∼70 dph. Birds were implanted with fiber optics when they were >100 dph. Birds were given at least 1 week to recover from cannula implantation and to habituate to singing with attached optical fibers. Custom LabView software (National Instruments) was used for online detection of preselected syllable templates and implementation of closed-loop optogenetic manipulation. 100-ms light pulses were delivered over a subset of variants of the target syllables in real time (system delay less than 25ms) for 3-12 consecutive days. 1.5-4mW of ∼520nm or ∼460nm LED output was delivered from the probe (200 or 250um, NA=0.66, Prizmatix, Israel). Acoustic signals were recorded continuously by a microphone (Shure BETA 98A/C) immediately adjacent to the bird’s cage using Sound Analysis Pro2011(SAP2011)(*44*) and bandpass filtered between 0.3 and 10 kHz.

### Song similarity scoring

Zebra finch song can be classified into three levels of organization: syllables, which are individual song elements separated by short silent gaps >5ms in duration; motifs, which are stereotyped sequences of syllables; and song bouts, which are defined as periods of singing comprised of introductory elements followed by one or more repeats of the song motif with inter-motif intervals <500ms. An automated procedure(*45*) was used to compute both song acoustic and sequence similarity of 30-50 song bouts from indicated time points to 3-6 representative motifs (including introductory notes) selected from baseline. Similarity index is defined as the product of the acoustic similarity and sequence similarity score.

### Longitudinal tracking song change

Song recordings were processed with ASAP (Automated Song Analysis Pipeline; v0.3.3, available at GitHub), an open-source R package for longitudinal acoustic analysis of learned vocalizations. Separate workflows were used for juvenile song development and adult song manipulation experiments.

#### Juvenile song development analysis

To track song development across days post-hatch (dph), recordings were analyzed using template-based detection and alignment. An acoustically distinct immature syllable that was reliably present across the developmental period was selected from a representative recording and used to generate a spectral template within a defined frequency range. Template matching was then applied across all recordings using cross-correlation, yielding detection times for each occurrence of the anchor syllable. Within a user-defined temporal window, only the highest-confidence match was retained.

Song units were segmented in one of two ways depending on the analysis: (i) fixed windows surrounding anchor detections were used to extract stereotyped motifs, or (ii) continuous song bouts were identified by merging adjacent vocalizations separated by less than a minimum silence interval. Extracted segments were aligned to the first anchor detection within each unit, allowing homologous acoustic structure to be compared across recording days. For visualization of developmental trajectories, amplitude envelopes from hundreds of aligned song renditions were computed and displayed as heatmaps grouped by dph. Timestamps corresponding to the aligned song motifs were exported for subsequent alignment of fiber photometry signals to a common behavioral reference.

#### Adult song tracking during optical manipulation

To quantify song changes in adult birds during optogenetic manipulation, individual motifs were isolated using the same template-based anchor detection and fixed-window extraction procedure. For latent-space analysis, each motif was represented by a time-resolved spectrogram obtained with a sliding window, generating a high-dimensional acoustic feature matrix across motif time points. To avoid sampling bias, equal numbers of motifs were drawn from each experimental epoch. The resulting feature matrix was reduced by principal component analysis (PCA), and the leading components were embedded in two dimensions using Uniform Manifold Approximation and Projection (UMAP)(*46*). Each motif rendition therefore formed a continuous trajectory through the shared latent space. Motif trajectories from Pre and Rec periods were projected into the same embedding for visual comparison of song structure before and after optical manipulation.

### Quantification of inhibition windows

Inhibition windows relative to motif onset or offset were estimated from 30 song bouts per bird and subsequently used for information theory analysis. To reduce the influence of other factors, such as motif duration or silence-gap duration, the temporal distance was defined as the time between the offset of the light pulse and either motif onset or motif offset.

For light pulses that overlapped with two consecutive motifs, the following motif was used for quantification. To compare the temporal relationship between optical inhibition and motif timing in DEC and STB birds, we calculated the average distance between light-pulse offset and motif onset for each bird. For birds producing more than one motif per song bout, we used the distance between light-pulse offset and the onset of the immediately following motif whenever that motif occurred within the same song bout, resulting in negative values when the light pulse ended before motif onset. Otherwise, the motif overlapping with the light pulse was used for quantification, resulting in positive values. We then used a nonparametric bootstrap approach to estimate the distribution of temporal distances for DEC and STB birds. Specifically, observed values within each group were randomly sampled with replacement 1,000 times to generate bootstrap samples of the same size, from which 95% CIs were computed.

### Quantification of song features

50 random selected song bouts per timepoint were used to estimate the latency for song initiation, omitted and repeated vocal elements in individual birds. The duration from the first vocal element of a song bout to the first syllable was defined as the latency for song initiation. Probability of individual syllables in a song bout was computed to demonstrate the frequency of syllable omission at indicated timepoints. The number of repetitions of introductory notes prior to the first motif was quantified. Song bouts generated within the first two hours on the Pre, last Post, or Rec days were analyzed to estimate song production motivation.

### Variable classification for DEC and STB assignment

A random forest model was used to evaluate how strongly individual song and inhibition-related factors contributed to distinguishing DEC birds from STB birds(*47*). For each predictor, variable importance was calculated using the varImp function(*48*). In this context, “importance” reflects how much a given variable improves classification performance across the decision trees in the forest. Variables with higher importance scores contribute more to reducing classification error or improving node separation when the model assigns birds to the DEC or STB group.

Mutual information (*I*) was then computed to measure how much knowing an individual factor, X, reduces uncertainty about the group identity, Y, where Y represents whether a bird was classified as DEC or STB. The mutual information between a given factor, X, and group identity, Y, can be expressed as:

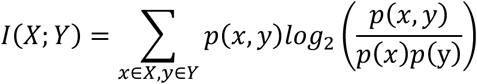

The mutual information between a joint variable constructed from two factors, X1 and X2, and bird group identity, Y, can be expressed as:

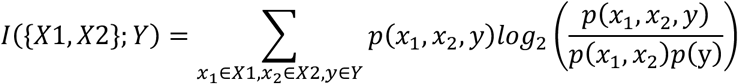

A quantile-based algorithm was used to discretize each individual factor, X, into four bins, with each bin containing a similar number of birds. Here, *p*(*x*) represents the probability that a bird falls within a given bin for a specific factor, and *p*(y) represents the probability that a bird belongs to either the DEC or STB group. The joint probability *p*(*x*, *y*) represents the probability that a bird belongs to a specific factor bin and a specific group. Mutual information was calculated using the Python package dit(*49*).

To assess the significance of mutual information for each factor, we generated 1,000 surrogate datasets by shuffling the observed group labels while preserving the bin assignments for each factor. Information-theoretic analyses were then performed on these surrogate datasets to generate a null distribution of mutual information values. The p-value for each factor was estimated as the proportion of null values greater than or equal to the observed mutual information.

### Statistic

The Shapiro-Wilk test was performed for all behavioral data to test for normality of underlying distributions. Repeated measure and ordinary one-way ANOVA followed by Bonferroni multiple comparison tests were used where appropriate.

If data were not normally distributed, non-parametric two-tailed testing was performed. Mann-Whitney tests and Mood’s tests were used where appropriate. Kruskal-Wallis tests were performed when comparisons were made across more than two time points from individual animals, whereas Friedman tests were performed when data was pooled across animals and comparisons were made across more than two time points. Statistical hypothesis testing was done at a 0.05 significance level. Statistical significance is represented as *p < 0.05, **p < 0.01, ***p<0.001. Statistical analysis was performed using R Studio (2002.12.10+353), and Prism (GraphPad Software).

## Data availability

Processed photometry datasets are available through the UT Southwestern Research Data Repository. (https://dataverse.tdl.org/previewurl.xhtml?token=49f8ad9a-31e8-4964-b215-21c204f0688c). Source data are provided with this paper.

## Code availability

R and Python code for reproducing Fig. 1 and 3 is available at https://git.biohpc.swmed.edu/xlei/da2song. Code for reproducing Fig. 2 is available at https://lxiao06.github.io/Juvenile_DA_analysis/. Original functions for song analysis are available at https://github.com/LXiao06/ASAP. Functions for photometry data analysis and TD learning-related metrics are available at https://github.com/LXiao06/VocalNeuroSync.

## Declaration of generative AI and AI-assisted technologies

During the preparation of this manuscript, the authors used Anthropic Claude Opus 4.7 and OpenAI 5.4 to support the revision of functions and to provide feedback on drafts of selected sections. All resulting content was subsequently reviewed, edited, and validated by the authors, who assume full responsibility for the final version of the manuscript.

## Acknowledgements

We are grateful to Drs. Brenton Cooper, William Dauer, Daisuke Hattori, Attila Losonczy, Steven Shabel, and Wen-hao Zhang, as well as Jeremy Brusch and Erfan Zabeh, and members of the Roberts Lab for their valuable comments on earlier versions of this manuscript. We thank the laboratories of John Pearson and Richard Mooney for making data from the Qi et al. 2025 study publicly available. We thank Dr. Peng Lian for support with the research container and the BioHPC supercomputing facility (Lyda Hill Department of Bioinformatics, UT Southwestern Medical Center) for providing computational resources. We also thank Dr. Massimo Trusel, Rahulraj Mishra, and Ziran Zhao for their assistance with surgery, optogenetic, and photometry experiments, and Dr. Guangzhong Wang and Mou Cao for their contributions to ASAP development, Jennifer Holdway, Luis Garcia, and Rico Cabuco for laboratory support. We thank Jesse Hilton, Richonda Hunte, and Pamela Jennings for administrative support. We thank the National Institutes of Health for supporting this research through R01 NS102488 to TFR.

## Author Contributions

LX and TFR conceived the project. LX designed the methodology and performed all experiments. LX and TFR visualized the project. TFR acquired funding. TFR administered and supervised the project. LX wrote the original draft of the manuscript. LX and TFR contributed to writing, reviewing and editing the manuscript. All data is available in the main text or the supplementary materials and links.

## Additional Information statement

Supplementary Information is available for this paper.

Correspondence and requests for materials should be addressed to TFR

